# Debugging and consolidating multiple synthetic chromosomes reveals combinatorial genetic interactions

**DOI:** 10.1101/2022.04.11.486913

**Authors:** Yu Zhao, Camila Coelho, Amanda L. Hughes, Luciana Lazar-Stefanita, Sandy Yang, Aaron N. Brooks, Roy S. K. Walker, Weimin Zhang, Stephanie Lauer, Cindy Hernandez, Leslie A. Mitchell, Neta Agmon, Yue Shen, Joseph Sall, Viola Fanfani, Anavi Jalan, Jordan Rivera, Feng-Xia Liang, Giovanni Stracquadanio, Lars M. Steinmetz, Yizhi Patrick Cai, Jef D. Boeke

## Abstract

The Sc2.0 project is building a eukaryotic synthetic genome from scratch, incorporating thousands of designer features. A major milestone has been achieved with the assembly of all individual Sc2.0 chromosomes. Here, we describe the consolidation of multiple synthetic chromosomes using endoreduplication intercross to generate a strain with 6.5 synthetic chromosomes. Genome-wide chromosome conformation capture and long-read direct RNA sequencing were performed on this strain to evaluate the effects of designer modifications, such as loxPsym site insertion, tRNA relocation, and intron deletion, on 3D chromosome organization and transcript isoform profiles. To precisely map “bugs”, we developed a method, CRISPR Directed Biallelic *URA3*-assisted Genome Scan, or “CRISPR D-BUGS”, exploiting directed mitotic recombination in heterozygous diploids. Using this method, we first fine-mapped a *synII* defect resulting from two loxPsym sites in the 3’ UTR of *SHM1*. This approach was also used to map a combinatorial bug associated with *synIII* and *synX*, revealing a highly unexpected genetic interaction that links transcriptional regulation, inositol metabolism and tRNA_Ser_^CGA^ abundance. “Starvation” for tRNA_Ser_^CGA^ leads to insufficient levels of the key positive inositol biosynthesis regulator, *Swi3*, which contains tandem UCG codons. Finally, to expedite consolidation, we employed a new method, chromosome swapping, to incorporate the largest chromosome (*synIV*), thereby consolidating more than half of the Sc2.0 genome in a single strain.

## Introduction

Rapid advances in DNA synthesis technology enable the possibility of transitioning from genome reading and editing to genome writing. We are designing and synthesizing a eukaryotic genome *in silico* through a bottom-up approach (Richardson et al., 2017). This Sc2.0 genome is based on that of *Saccharomyces cerevisiae*, a unicellular eukaryotic model organism widely used in basic research and industrial fermentation. Previously, 6.5 out of 16 chromosomes had been successfully synthesized (Annaluru et al., 2014; Dymond et al., 2011; Mitchell et al., 2017; Shen et al., 2017; Wu et al., 2017; Xie et al., 2017; Zhang et al., 2017). Since then, the assembly of all the synthetic chromosomes has been completed. Each synthetic chromosome was synthesized separately by teams comprising the international Sc2.0 consortium. Consequently, each yeast strain produced contains only one synthetic chromosome, with the remainder of the genome still native. A challenge is to consolidate every chromosome into one fully synthetic Sc2.0 strain. In this study, we used an “endoreduplication intercross” strategy to consolidate all previously constructed 6.5 synthetic chromosomes (*synII, synIII, synV, synVI, synIXR, synX* and *synXII*) into a single strain.

The design of Sc2.0 will boast thousands of genome-wide edits as designer features. One of these is the relocation of all tRNAs to a neochromosome, requiring the removal of all endogenous tRNAs. Native *S. cerevisiae* contains 275 genomic tRNAs. In the syn6.5 strains, 97 of these were deleted, accounting for one-third of tRNA pools. To avoid possible growth defects due to reduced abundance of tRNAs during consolidation, we integrated a tRNA array into each synthetic chromosome, consisting of the synthetic counterparts of all the tRNAs deleted from their original locus on that chromosome. Meanwhile, other extensive modifications to the genome can also result in unexpected bugs in the form of unwanted fitness defects. Precisely and systematically mapping these variants has been a challenging and laborious task. Inspired by diverse CRISPR applications in yeast genomic editing, regulation and mapping (DiCarlo et al., 2013; Jacobs et al., 2014; Sadhu et al., 2016; Zetsche et al., 2015; Zhao and Boeke, 2018; Zhao and Boeke, 2020), we developed a highly reliable bug mapping method known as CRISPR Directed Biallelic *URA3*-assisted Genome Scan (CRISPR D-BUGS). We successfully repaired bugs identified in single synthetic chromosomes, including a mitochondria-related defect caused by two loxPsym sites in the 3’ UTR of *SHM1*. CRISPR D-BUGS was also expanded to map a combinatorial defect associated with an essential tRNA gene *SUP61* in *synIII* and *SWI3* in *synX*, encoding a subunit of the SWI/SNF chromatin remodeling complex. In this case, neither variant alone caused a fitness defect, but the combination of the two caused a severe defect.

In strains with multiple synthetic chromosomes, all mobile elements were deleted, tRNAs were relocated, and loxPsym sites were inserted into the 3’ UTRs of many genes by design. To probe the effects of these changes on the 3D genome organization and transcriptional regulation of the compact yeast genome, we used chromosome conformation capture (Hi-C) and Nanopore direct RNA sequencing (Garalde et al., 2018) to characterize a strain with 6.5 synthetic chromosomes.

Finally, to expedite consolidation, we used a new method, chromosome swapping, to transfer the largest single synthetic chromosome, *synIV*, into the yeast strain that already carried 6.5 synthetic chromosomes, thereby consolidating more than half of the genome of Sc2.0, and producing the syn7.5 strain.

## Results

### Synthetic chromosome consolidation using endoreduplication intercross

The Sc2.0 consortium assembled each of the 16 synthetic chromosomes (*synI-synXVI*) in discrete haploid strains. Two strains of opposite mating types and carrying different synthetic chromosomes can mate to generate a heterozygous diploid strain with multiple synthetic chromosomes. In order to avoid generating chimeric chromosomes due to efficient meiotic crossovers during sporulation, we previously established a consolidation strategy called “endoreduplication intercross”, which takes advantage of inducible chromosome destabilization (Mitchell *et al*., 2017; Reid et al., 2008). Briefly, two strains with different synthetic chromosomes and opposite mating type are mated; the resulting heterozygous diploid strain carries two synthetic chromosomes along with their native counterparts, and the latter can be specifically destabilized using a pGAL promoter inserted adjacent to its centromere. After sporulating and screening the resulting spore clones, we obtain haploid strains with two or more synthetic chromosomes. This strategy was used to sequentially combine more than two individual synthetic chromosomes from their discrete parental strains. Following several rounds of intercross consolidation in which one new synthetic chromosome is consolidated per round, we obtained a single strain, YZY1178, with 6.5 synthetic chromosomes (*synII, synIII, synV, synVI, synIXR, synX* and *synXII)*, representing all synthetic chromosomes assembled previously (Figure 1A and S1). In this strain, ∼31% of the genome is synthetic, and thus it carries many designer features, including 91 removed introns, 97 relocated tRNAs, 444 TAG stop codons swapped to TAA, more than one thousand inserted loxPsym sites, and 4814 pairs of synonymously recoded synthetic PCRtags (Figure 1B).

**Figure 1.**
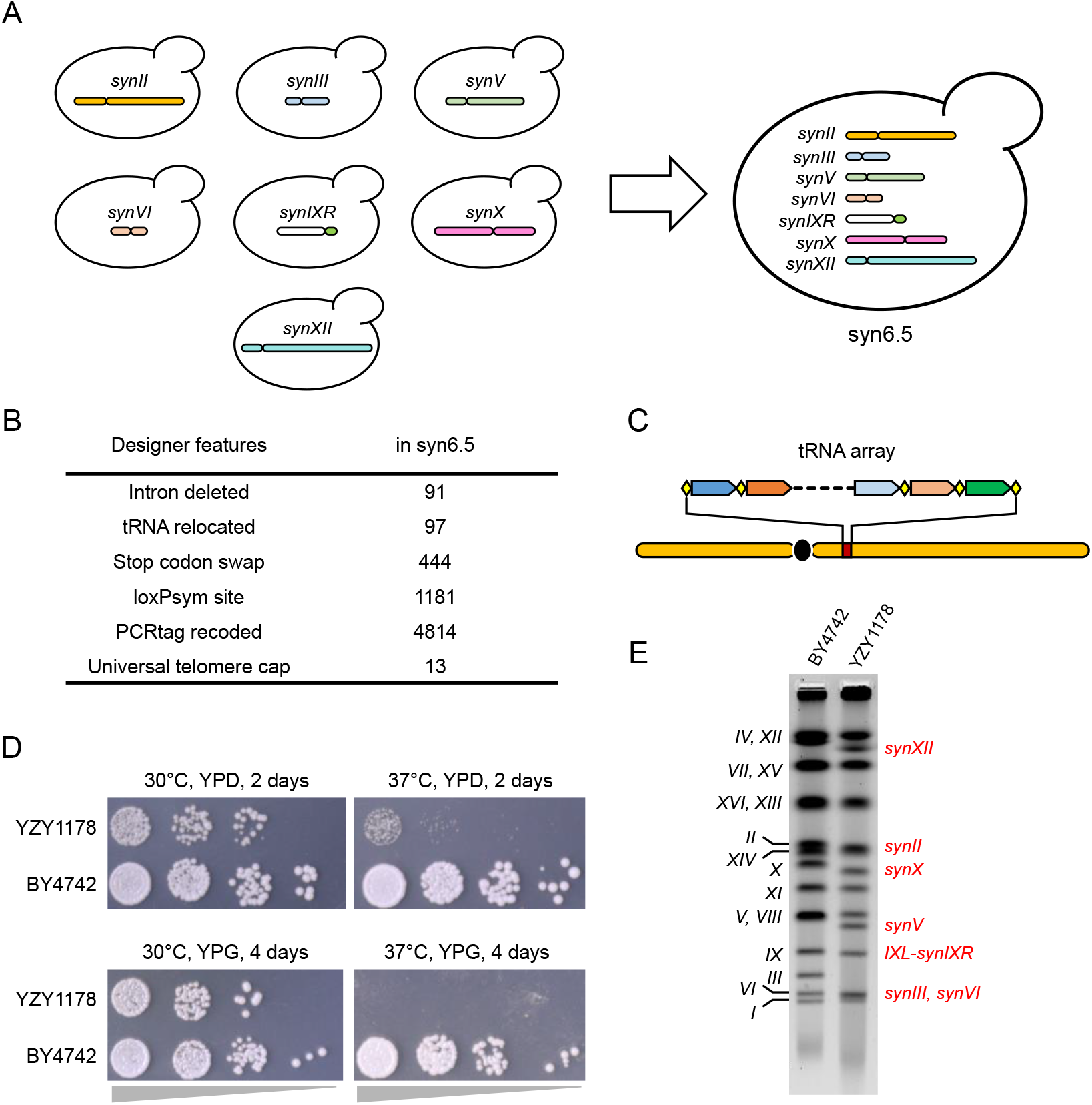
Consolidation of multiple synthetic chromosomes. (A) All previously assembled synthetic chromosomes, *synII, synIII, synV, synVI, synIXR, synX* and *synXII* were consolidated using endoreduplication intercross, generating one haploid strain, syn6.5. (B) Sc2.0 designer features carried in the syn6.5 strain. (C) A tRNA array was integrated into each synthetic chromosome to maintain the tRNA abundance and balance. Each tRNA gene was flanked with rox recombination sites (yellow diamond). The detailed anatomy of these arrays is shown in Figures S3 and S4. (D) Fitness assays for draft syn6.5 strain, YZY1178, after consolidation was completed. (E) Pulsed field gel (PFGE) to evaluate the electrophoretic karyotype of a syn6.5 strain.

One design feature of Sc2.0 is removal of all tRNA genes from their native loci, for eventual relocation to a specialized tRNA neochromosome (Richardson *et al*., 2017). This feature creates a practical challenge for consolidating the chromosomes - before neochromosome assembly and delivery is complete, the available number of tRNA genes will decrease as more and more synthetic chromosomes are consolidated in a haploid strain. Notably, the tRNA genes lacking from the 6.5 synthetic chromosomes represent 97 of the 275 genomic tRNAs (Table S1). As an interim solution for possible fitness defects caused by this tRNA deficit, we designed a tRNA array, consisting of all tRNA genes from each individual chromosome, and then integrated each of these into its host synthetic chromosome (Figure 1C), thus maintaining the tRNA abundance and balance as additional synthetic chromosomes are incorporated. Briefly, each tRNA array was released from its host plasmid and integrated using homologous recombination (Figure S2). All tRNA genes were also flanked by rox sites, which can be recognized by Dre (but not Cre) recombinase, enabling a chromosomal tRNA-specific rearrangement system (Figure S3-S4).

The “draft” syn6.5 strain grows slightly slower on rich medium (YPD) but grows comparably on plates with non-fermentable glycerol (YPG) (Figure 1D). Unlike the parent strains, the syn6.5 strain also shows an obvious growth defect at high temperature (37°C), suggesting the existence of a new “combinatorial bug” resulting from genetic interactions between designer variants introduced on more than one synthetic chromosome, analogous to the phenomenon of synthetic lethality/fitness. The karyotype of this strain was confirmed using pulsed field gel analysis (PFGE), with synthetic chromosomes showing expected faster migration due to their shorter lengths (Figure 1E). The uniform genome coverage of each chromosome in whole genome sequencing (WGS) (Figure S5). No new mutations or genome arrangements appeared during consolidation when compared to the original parental strains with single synthetic chromosomes.

### Mapping fitness defects using CRISPR D-BUGS

With thousands of designer modifications introduced, growth defects or “bugs” resulting from some of these designer changes have been observed in most synthetic chromosomes. Identifying these bugs is important for repairing cell fitness,and understanding their mechanisms may illuminate new biological insights. We developed a systematic and efficient bug-mapping strategy that exploits loss of heterozygosity (LOH) in diploids, called CRISPR Directed Biallelic *URA3*-assisted Genome Scan, or CRISPR D-BUGS. This approach makes use of heterozygous diploid strains bearing a synthetic chromosome and a native chromosome in which one of the telomeres bears a *URA3* marker gene. In such diploid strains, homologous recombination between two chromatids can be enhanced by a targeted chromosomal double-strand break (Figure 2A) (Sadhu *et al*., 2016). After cell division, daughter cells will carry a pair of chromosomes that are homozygous for synthetic DNA from the recombination site to the telomere region but retain heterozygosity in the remainder of the chromosome; these LOH events can be readily selected for by plating on 5-FOA. By using gRNAs that target different PCRTag sequences, a series of yeast strains with various homozygous synthetic regions can be generated (Figure 2B). We checked the fitness of the strains and subsequently mapped the “fitness boundary” at which derivative strains shift phenotypically from unhealthy to healthy.

**Figure 2.**
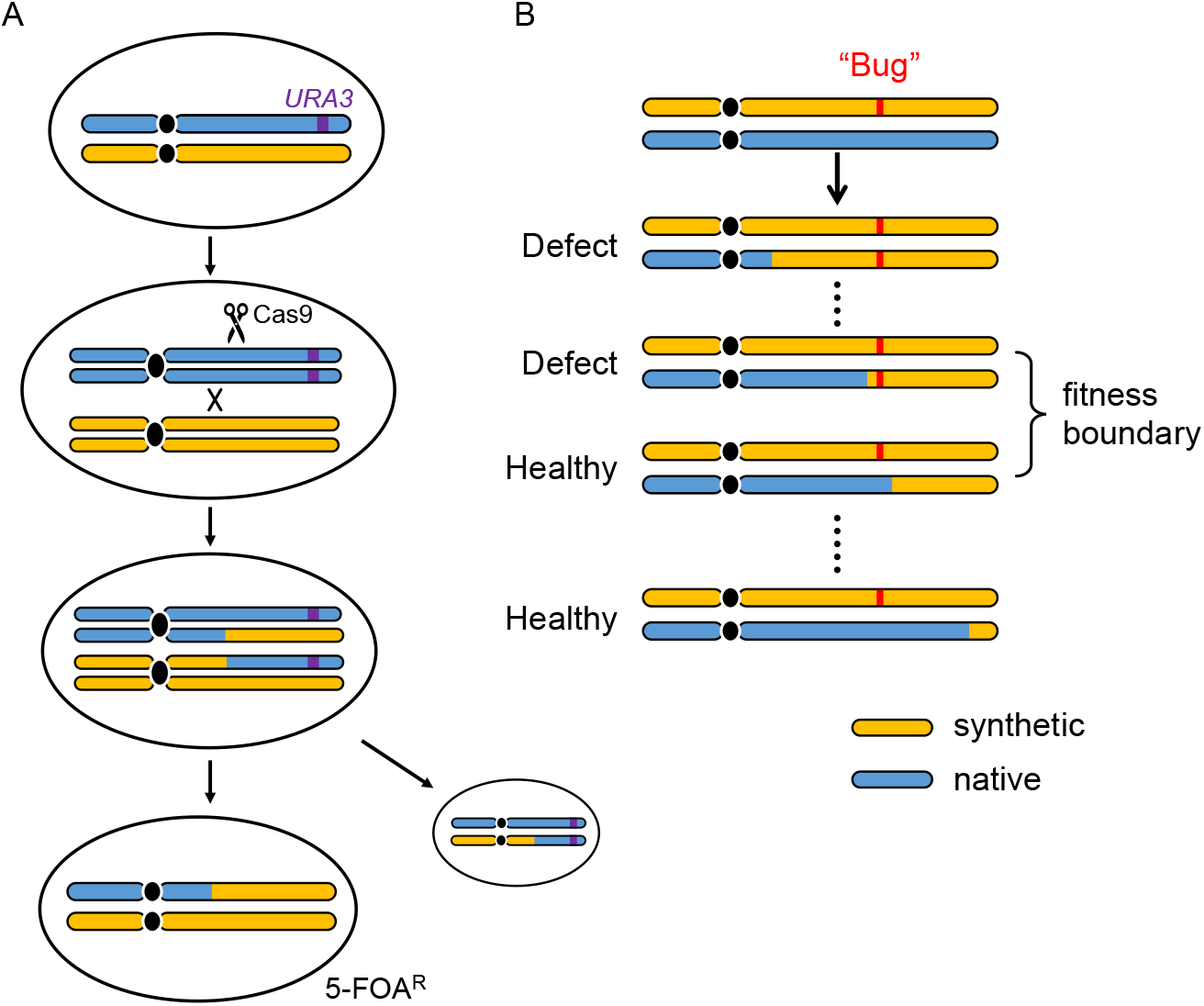
Fitness mapping using CRISPR D-BUGS. (A) General outline. A *URA3* marker is integrated into the native allele (blue), which is cleaved by Cas9 targeted by an sgRNA selected specifically to cut at one WT PCRtag. Following mitotic recombination, strains homozygous for the synthetic region (orange) are selected on 5-FOA plate. (B) A series of these strains with different synthetic region are generated by gRNAs targeting different WT PCRtag loci to map the fitness boundary.

Absent other mapping information, screening is begun within the resolution of a chromosome arm. Subsequently, the search continues with a series of gRNAs to map the bug more precisely. Resolution can be increased with two or more rounds of mapping until a group of colonies generated from the same single gRNA shows mixed fitness. This variability results from mitotic recombination occurring within a ∼10kb window from the DNA cleavage site, such a region may include multiple designer features, such as PCRtag or loxPsym insertions (Sadhu *et al*., 2016). Using WGS, the bug is mapped at high resolution by defining the locations of the synthetic genome modifications within the recombination interval. Following similar principles, CRISPR D-BUGS can also be used to map dominant bugs (Figure S6).

To test this approach, we first tried to map a perplexing bug on a previously synthesized chromosome. The original *synII* strain (chr02_9_03) showed a growth defect on YPG medium at 37°C (Figure 3A). This recessive defect appeared after megachunk X was assembled (Shen *et al*., 2017). To map the *synII* bug using CRISPR D-BUGS, we constructed a *synII*/+ heterozygote, and then selected gRNAs targeting PCRtags within megachunk X. The colonies generated from gRNA.YBR256C and gRNA.YBR261C all showed the defect, whereas colonies generated from gRNA.YBR270C and gRNA.YBR275C were all healthy (Figure 3B and S7). This mapped the bug between *YBR261C* and *YBR270C*. In a second round of bug mapping, single colonies generated using gRNA.YBR265W showed a mixture of two fitness levels (Figure 3B and S8). Using WGS, we precisely located the recombination interval of each colony based on the synthetic sequence variants, and linked each variant to strain fitness (Figure 3C). This strategy helped identify two adjacent loxPsym sites between *YBR263W* (*SHM1*) and *YBR264C* (*YPT10*) as responsible for the fitness defect. Deleting both loxPsym sites successfully restored strain fitness, as in strain YZY166 (chr02_9_04) (Figure 3A).

**Figure 3.**
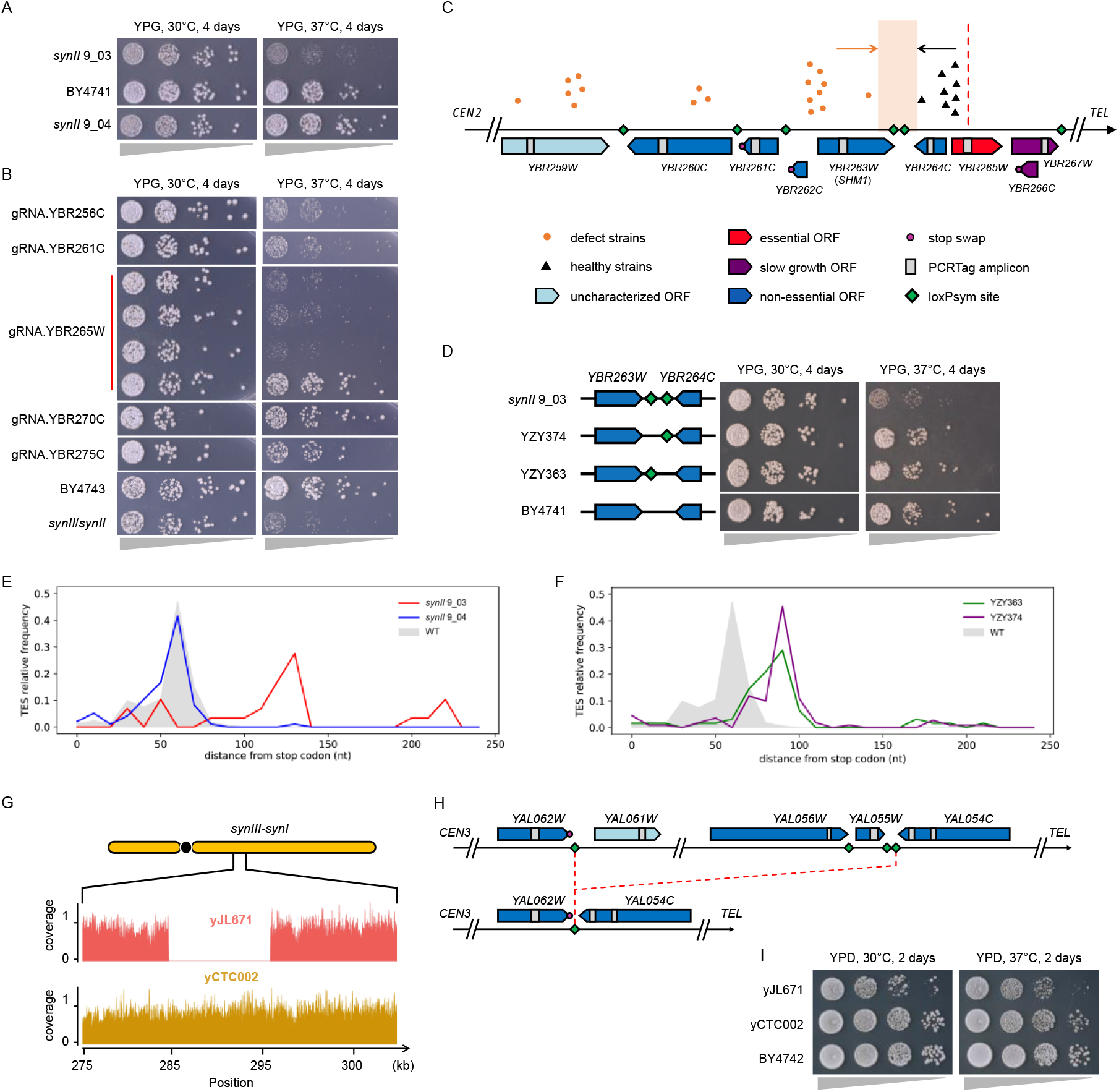
CRISPR-D-BUGS mapping in *synII* and *synIII-synI* fusion chromosomes. (A) Fitness assay on YPG plates for a strain with the original *synII* (9_03), compared to wild type control (BY4741). In strain YZY166 (9_04), the bug was fixed by deleting the two loxPsym sites downstream of *SHM1*. (B) The CRISPR-D-BUGS colonies generated using gRNAs labeled on the left side. For each gRNA, at least four colonies were tested and showed the same fitness except gRNA.YBR265W. More colonies are shown in Figure S7. (C) The recombination sites in gRNA.YBR265W colonies indicating their fitness level are aligned with *synII* designer features. Red dashed line indicates the locus in *synII* corresponding to the cleavage site in the native counterpart. The original fitness assay for these strains is shown in Figure S8. (D) Fitness assay for the strains with the deletion of both or either loxPsym sites(s). (E) Transcript end site (TES) distributions of *SHM1* transcripts from original *synII* (red) and updated *synII* (blue), compared to wild type (gray). (F) The same measurements in strains with either loxPsym site deleted (YZY363 and YZY374 as in panel D) are also shown. (G) Deletion detected in *synIII-synI* strain (yJL671), which was repaired in the final version (yCTC002). (H) The diagram of *synI* in design (upper) and actual strain of yJL671 (bottom). (I) Fitness assay for final *synI* strain (yCTC002) after bug was repaired.

The loxPsym sites were integrated 3bp downstream of the stop codon of *SHM1* and *YPT10*, a pair of convergent and closely spaced genes (Figure S9A). *SHM1* encodes the mitochondrial serine hydroxymethyltransferase, and its deletion results in impaired respiratory function, consistent with the observed *synII* fitness defect on YPG (May et al., 2020). In contrast, *ypt10* deletion showed no difference compared to wild-type strains under various conditions including different temperatures or carbon sources (Louvet et al., 1999). These genes are convergent and closely spaced, such that integration of loxPsym sites 3 bp downstream of their stop codons produces *SHM1* transcripts containing two loxPsym sequences in their 3’ UTRs. These two loxPsym sequences are predicted to form a stem-loop structure in the *SHM1* 3’ UTR, which we hypothesize may affect mRNA stability (Figure S9B and C). Consistent with this hypothesis, deletion of both or either loxPsym site(s) significantly recovered transcript abundance and successfully rescued the growth defect, strongly implying that formation of a stem loop in the RNA leads to loss of RNA abundance and the fitness defect (Figure 3D and S10A). We also found that *Shm1p* level was reduced in the presence of two loxPsym sites, and recovered upon their removal (Figure S10B).

Since the two loxPsym sites are located in the 3’ UTR, we also wondered about their effects on transcript properties. To answer this, we used Nanopore direct RNA sequencing to evaluate full-length native transcripts of *SHM1* and *YPT10* directly, and measured the transcript end site (TES) distributions (Brooks et al., 2022). In the original *synII* (chr02_9_03) strain, the majority of *SHM1* TESs were extended by ∼66 nt, matching the length of the two transcribed loxPsym sites (68 nt), indicating that *SHM1* transcript termini were not significantly affected (Figure 3E). There were also around 10% transcripts extended by ∼160 nt, forming a second peak specific to *synII*, suggesting that transcription termination could be slightly affected by the loxPsym sequences. The removal of both loxPsym sites (chr02_9_04) successfully recovered the TES distribution, overlapping with the wild type peak. For the *SHM1* TES in YZY363 and YZY374, in which the individual loxPsym sites were deleted, a single peak was formed and extended by ∼32 nt, matching the expected length of a single loxPsym site. We also measured the *YPT10* TESs in these strains and observed similar patterns (Figure S11). In summary, the incorporation of two loxPsym sites, presumably forming a stem loop in their 3’ UTRs, mainly affected the quantity but not the isoform boundaries of *SHM1* and *YPT10* transcripts.

### Mapping a *synI* bug to an unexpected deletion

CRISPR D-BUGS was also applied to map the growth defect of a *synI* strain. A special feature of *synI* is that it is fused with *synIII* (Luo et al., 2022). The draft strain (yJL671, chr01_9_02) showed a recessive growth defect even on rich medium (YPD), which is not caused by chromosome fusion (Figure S12). Using CRISPR D-BUGS, we successfully mapped the bug to a window of 5 open reading frames (ORFs) between *YAL055W* and *YAL049C* in a single round of mapping (Figure S13). Using WGS, we found an unexpected ∼10kb deletion in yJL671, which became homozygously deleted in the low-fitness diploids, but remained heterozygous in the healthy strains (Figure 3G and S14). By checking *synI*’s assembly history, we found that this deletion was caused by an unexpected off-target recombination between two loxPsym sites during CRISPR-mediated repair of a missense mutation in strain yJL663, which contained an earlier draft *synI* version (Figure 3H and S15A). By repairing this deletion using SpCas9-NG, a final healthy strain with the complete *synI* sequence was obtained as yCTC002 (Figure 3I and S15B) (Nishimasu et al., 2018). In summary, CRISPR D-BUGS was used to quickly map distinct bug types in *synI* and *synII*.

### A combinatorial bug associated with *synIII* and *synX*

Although strains with single synthetic chromosomes are healthy, combinatorial defects can still occur due to combinations of sequence changes that by themselves have no phenotypes, owing to genetic interactions between variants on two (or theoretically more than two) synthetic chromosomes. While strains containing *synIII* (chr03_9_02) and *synX* (chr10_9_01) alone are healthy, we found a combinatorial defect at high temperature in a strain containing both *synIII* and *synX* and no other synthetic chromosomes (Figure 4A). We first used CRISPR D-BUGS to map the bug in *synX* (Figure S16). In the first round of “rough” mapping, we successfully mapped a fitness boundary between *YJL097W* and *YJL210W* at the left arm (Figure S17). By fine mapping within this interval, we found that single colonies generated from a gRNA targeting *YJL176C* showed mixed fitness levels (Figure. S18). Using WGS, we mapped the bug to the loxPsym site integrated downstream of the *YJL175W* ORF, representing the boundary separating healthy and temperature-sensitive strains (Figure 4B).

**Figure 4.**
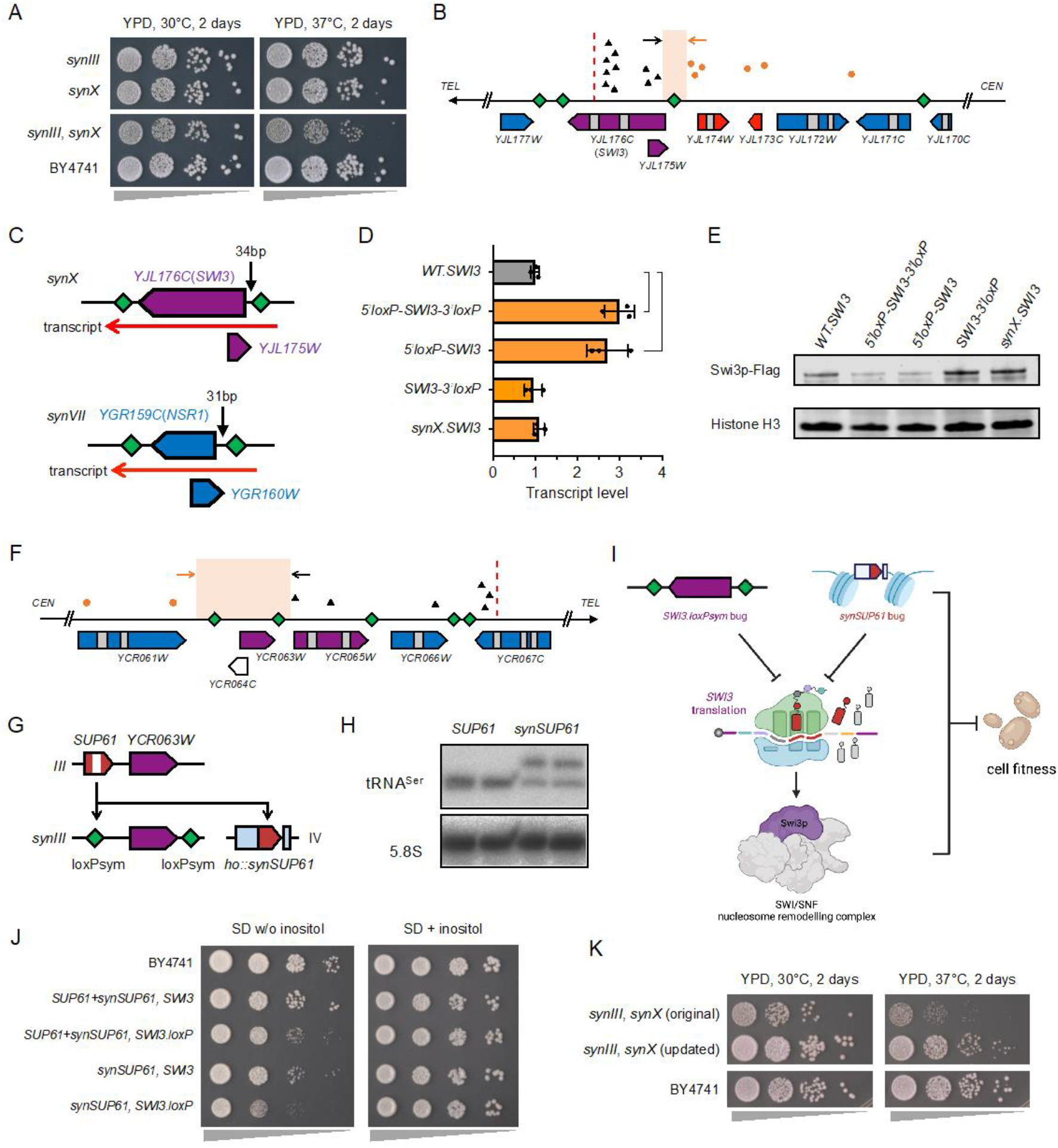
Combinatorial defect between *synIII* and *synX*. (A) Fitness assay showing a combinatorial defect in *synIII, synX* context. (B) For *synX* bug mapping, single colonies generated using gRNA.YJL176C showed mixed fitness; recombination intervals were aligned to *synX* left arm. Same labels as in Figure 3C were used here. Fitness assays for these colonies are shown in Figure S18. (C) Diagram of *YJL176C* (*SWI3*) loxPsym pattern, compared to *YGR159C* (*NSR1*) from *synVII* (Shen et al., 2022). (D) *SWI3* transcript levels in wild type background (gray bar), and *synX* strains (orange bars) with both loxPsym sites (*5’loxP-SWI3-3’loxP*), 3’ loxPsym deleted (*5’loxp-SWI3*), 5’ loxPsym deleted (*SWI3-3’loxP*) and no loxPsym site (*synX*.*SWI3*). (E) Immunoblotting of Swi3p-Flag in strains with the loxPsym deleted from 5’ and/or 3’ UTR. (F) For *synIII* bug mapping, recombination intervals of single colonies generated from gRNA.YCR067C were aligned to *synIII* right arm. Fitness assays for these colonies are shown in Figure S20. (G) Relocation of *synSUP61* to the *HO* locus. Gray blocks indicate flanking sequences from *Ashbya gossypii*. White band indicates the *SUP61* intron that is removed in the synthetic version. (H) Northern blot to check the quality and level of tRNA^Ser^ expressed from native and synthetic *SUP61*. (I) Proposed combinatorial interactions between *synSUP61* bug and *SWI3*.*loxP* bug. (J) Inositol auxotrophy analysis, with wild type *SUP61* integrated and/or *SWI3* bug repaired. (K) Fitness assay of final *synIII, synX* strain with both bugs fixed.

*YJL175W* is a “dubious ORF”, and overlaps the 5’ end of *YJL176C* (*SWI3*), an important named gene. The loxPsym inserted into the 3’ UTR of *YJL175W* is transcribed as a part of the *SWI3* 5’ UTR (Figure 4C). Consequently, there are two loxPsym sequences in the transcript of *SWI3*, which might therefore form a looped secondary RNA structure. Interestingly, the *synX SWI3* transcript level was increased by three-fold, and this transcriptional phenotype was restored to the wild-type level by deleting the 5’ UTR loxPsym, but not by deleting the 3’ UTR loxPsym (Figure 4D). Paradoxically, the Swi3p level was reduced in the presence of 5’ UTR loxPsym and restored upon its removal (Figure 4E). The most parsimonious explanation for these results is that the inverted repeat within the loxPsym sequence in the 5’ UTR forms a stem loop that stabilizes the RNA isoform and blocks translation. Insertion of the loxPsym in the 3’ UTR, as in all other nonessential Sc2.0 genes, had minimal to no effect on protein levels. The temperature sensitive phenotype unique to the *synIII, synX* strain (but not the two parental strains) is consistent with the fact that a *swi3* null allele results in temperature sensitivity (Auesukaree et al., 2009). As an essential component of the SWI/SNF chromatin remodelling complex, Swi3p is required for the transcription of many genes, including *INO1*, and *swi3* null mutants are viable but auxotrophic for inositol (Peterson and Herskowitz, 1992; Villa-García et al., 2011; Yoshinaga et al., 1992). Remarkably, the *synX* strain displayed partial inositol auxotrophy, which was largely restored by removing the 5’ UTR loxPsym site (but not the 3’ UTR loxPsym site; Figure S19), consistent with the proposed mechanism. A similar pattern was observed in a *synVII* bug (Figure 4C), wherein a similarly “misplaced” loxPsym site in the *NSR1* 5’ UTR led to increased mRNA, but dramatically reduced protein level (Shen et al., 2022). As in the case above, deletion of the 5’ loxPsym site restored normal transcript and protein levels, and rescued the growth defect in *synVII* strains.

Following the same principles, we mapped the bug in *synIII* (chr03_9_02) with CRISPR D-BUGS initially to the right arm, then roughly to between *YCR057C* and *YCR067C* (Figure S20) and finally fine-mapped the bug to two loxPsym sites between *YCR061W* and *YCR065W* (Figure 4F). By restoring them to wild-type, we found that the left side loxPsym site between *YCR061W* and *YCR064C* seemingly caused the defect (Figure S21). This loxPsym site marked the deletion of *SUP61*, an essential single copy tRNA_Ser_^CGA^ gene, which decodes the rare UCG serine codon (Figure 4G). Importantly, unlike the examples mentioned above, this loxPsym site was not embedded inside any transcribed region, suggesting that it might be deletion of the tRNA itself that was responsible for the defect. As a part of the overarching Sc2.0 design, all tRNA genes are to be relocated to a new synthetic neochromosome encoding only tRNAs (Richardson *et al*., 2017). Before introduction of the complete tRNA neochromosome, all strains containing *synIII* have a synthetic version of *SUP61* (*synSUP61*) integrated in the *HO* locus on *chrIV* to temporarily provide its essential function. By itself, *synSUP61* suffices for cell survival and health (Annaluru et al., 2014). Like all of the synthetic tRNAs in Sc2.0, *synSUP61* is flanked by 500bp 5’ and 40bp 3’ of *Ashbya gossypii* tRNA flanking sequences and has a precise intron deletion. Introducing a single copy of *SUP61* in the strain complemented the defect (Figure S21), suggesting that *synSUP61*is too lowly expressed or otherwise incapable of providing full functionality. To test this hypothesis, we examined expression of *synSUP61* by Northern blotting and observed that it produces only about half the normal amount of mature tRNA, and a surprisingly large amount of 5’ pre-tRNA, suggesting inefficient processing of this tRNA, relative to *SUP61* (Figure 4H). This appears to be associated with replacing the tRNA 5’ flanking region with the sequence from *Ashbya* in *synSUP61*, and not the intron deletion or the 3’ flanking region swap (Figure S22).

As both parental *synIII* and *synX* strains are healthy, we conclude that the combinatorial bug results from an unexpected interaction between *synSUP61* and *SWI3*, both of which appear to be under-expressed relative to their native counterparts. The single copy essential *SUP61* gene produces the only tRNA decoding the rare UCG serine codon. Interestingly, the *SWI3* transcript, has above average usage of UCG for serine, and importantly, it includes two tandem UCG codons (Figure S23). Tandem rare codons can cause translational pausing or even arrest due to “starvation” for charged cognate tRNAs (Guimaraes et al., 2020; Kane, 1995; Wang et al., 2016). Based on this, we hypothesized that reduced abundance of tRNA_Ser_^CGA^ further reduces expression of *SWI3* below the already lower than normal level, caused by the ectopic loxPsym site in the 5’ UTR. This is predicted to result in an even lower level of functional SWI/SNF complex and the resulting pronounced growth defect (Figure 4I). To test this hypothesis, we repaired either or both bugs, and measured inositol auxotrophy (Figure 4J). Interestingly, either deletion of the *SWI3* loxPsym site or addition of *SUP61* individually were able to partially rescue auxotrophy, suggesting that ultimately, the observed phenotypes are the consequence of low *Swi3* protein. When the two “buggy” components were both restored to their native forms, the fitness of the strain was successfully rescued. To confirm this combinatorial interaction mechanism, we mutated either of the *SWI3* tandem serine codons from rare UCG to common UCU, with *SWI3* loxPsym site deleted (Figure S24). Consistently, strains with either mutation showed significantly improved growth on plates without inositol. Notably, the inositol auxotroph was still not completely rescued. Similar results were also observed in *synX* strain, in which removal of the 5’ *SWI3* loxP site significantly, but not completely rescued inositol auxotrophy, suggesting that other synthetic modifications may exist affecting inositol biosynthesis.

By repairing the *SWI3* and *synSUP61* bugs, the fitness of the *synIII, synX* strain was largely rescued at both 30°C and high temperature (Figure 4K). For multiple synthetic chromosomes (*synII, synIII, synV, synVI, synIXR, synX, synXII*), we repaired all known bugs, including the *SHM1* bug in *synII* and the combinatorial bug between *SWI3* in *synX* and *synSUP61* in *synIII*. As expected, the growth defect was dramatically improved, albeit with minor residual growth defects at high temperatures (Figure S25).

### Characterization of multiple synthetic chromosomes in syn6.5 strains

Even after repairing bugs in the synthetic chromosomes that affect growth phenotypes, we remained curious about how multiple synthetic chromosomes would affect genome organization and whether the large numbers of designer features would affect transcription. Therefore, we used genomic chromosome conformation capture (Hi-C) to investigate the organization of all 6.5 synthetic chromosomes in the nucleus. The Sc2.0 design improved mappability as a result of the deletion of repetitive regions, especially the Ty elements (Mercy et al., 2017). In our strain with 6.5 synthetic chromosomes, we calculated the spatial contact frequency and generated a heat map of genomic interactions (Figure 5A). Based on that, we generated a 3D map showing the trajectories of the synthetic and native yeast chromosomes (Figure 5B and S27). Similar to the wild type, all centromeres of synthetic and native chromosomes interacted *in trans* near the spindle pole body (SPB), as well as their telomeres clustered at the nuclear envelope (Taddei and Gasser, 2012). To detect differences in the internal folding of the chromatin, we calculated the decay of contact frequency as a function of the genomic distance genome-wide, and no substantial differences were observed (Figure 5C). These results indicated that Sc2.0 designer modifications have minor effects on the chromosomal organization, even when multiple synthetic chromosomes are combined.

**Figure 5.**
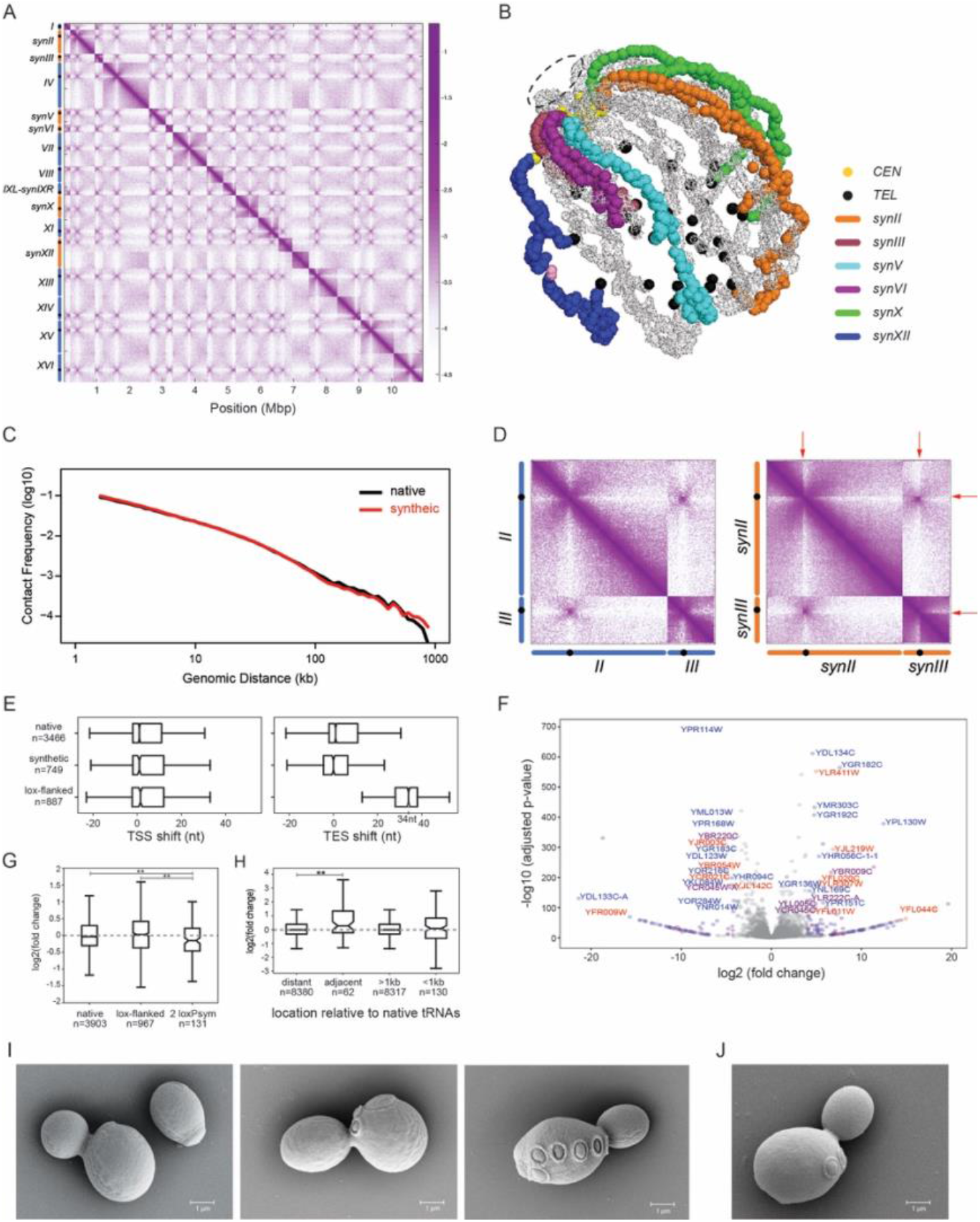
Characterization of the strain with multiple synthetic chromosomes. (A) The heat map shows the contact probability (log10) between pairs of chromosomal sites. The genome position (kb) and corresponding chromosomes were labeled as horizontal and vertical axes, respectively. (B) The 3D chromosome trajectories of multiple synthetic chromosomes. Gray, all other native chromosomes. The 3D structures are shown in Supplementary Data-1 and as a movie in Supplementary Data-2. (C) Contact frequencies as a function of the genomic distance. (D) Heat maps for native (left) and synthetic (right) chromosomes *II* and *III*. The sharp boundaries at the tRNA array integration loci are highlighted with red arrows. (E) Change in distribution of transcript start sites (TSSs) and transcript end sites (TESs) in transcript isoforms arising from the native and synthetic chromosomes in the syn6.5 strain compared to wild type. Lox-flanked genes indicate the genes on synthetic chromosomes with 3’ loxPsym sites. (F) Volcano plot of gene expression in the multiple synthetic chromosome strain, compared to wild type. Lists of significantly up- and down-regulated genes are presented in Tables S7 and S8. Blue, genes in native chromosomes. Purple, genes in synthetic chromosomes. Red, genes with loxPsym site incorporated. (G) Change in expression level of genes that give rise to transcripts with two loxPsym sites in their 3’ UTRs compared to genes on native and synthetic chromosomes in the syn6.5 strain. (H) Comparison of expression level changes in the synthetic strain, in which tRNAs are relocated, for the genes closest to native tRNAs (adjacent) and all other genes (distant), as well as genes either <1kb or >1kb away from native tRNAs. The adjacent genes are listed in Table S9. Mann Whitney U tests were used to determine significant difference between gene sets, **p<0.01. (I) SEM pictures of single yeast cells with multiple synthetic chromosomes, compared to wild-type cells as in (J).

Unexpectedly, for each synthetic chromosome, we noted a surprisingly sharp boundary formed at one position in their contact frequency maps (Figure 5D). These supersharp boundaries exactly match all locations of the tRNA arrays, such as the *synII* tRNA array integrated at the left arm close to *CEN2* (∼11 kb) and the *synIII* tRNA array integrated on the right arm close to *CEN3* (∼9.2 kb). The same results were observed in other synthetic chromosomes with tRNA arrays integrated, but not in the native chromosomes that lack these (Figure S26). This result suggests that very active tRNA transcription manifests as a higher frequency of intra-locus contacts for tRNA arrays and results in a mild structural alteration of the pericentromeric chromatin.

### Transcript profiling using RNAseq

To determine whether transcript boundaries were affected by the incorporation of synthetic design features, we mapped transcript isoforms from the syn6.5 strain using Nanopore long-read direct RNA sequencing. As expected, transcript start sites were not affected by inclusion of 3’ loxPsym sites. Neither transcripts arising from genes on the native chromosomes nor those without flanking loxPsym sites on the synthetic chromosomes showed end site alterations; however, the addition of loxPsym sites at the 3’ end of genes increased the length of their transcripts by 34 nt on average (Figure 5E). This is consistent with incorporation of the loxPsym site into the transcript without altering its cleavage/polyadenylation site.

To assess the effects of synthetic genome design on gene expression levels, we performed stranded mRNA sequencing. Some genes on the native and synthetic chromosomes, both with and without flanking loxPsym sites, showed significantly altered expression levels (Figure 5F), indicating that the synthetic genome design caused some, but not widespread, indirect effects on gene expression levels. LoxPsym-flanked genes were not significantly affected compared with genes on the native chromosomes; however, genes with transcripts that incorporate two loxPsym sites within their 3’ UTRs tended to experience a slight decrease in transcript abundance (Figure 5G), potentially indicating decreased stability of these transcripts on average. This was consistent with the *synII* growth defect caused by two tandem loxPsym sites in the *SHM1* 3’ UTR. The relocation of tRNAs led to a major alteration in the 3D organization of the synthetic chromosomes (Figure 5D). We therefore compared expression of genes adjacent to tRNAs in the native and synthetic chromosomes and saw that the removal of tRNA genes in the synthetic genome appeared to be associated with increased expression of their former neighbors (Figure 5H), consistent with previous studies of tRNA gene mediated silencing (Good et al., 2013; Hamdani et al., 2019; Hull et al., 1994). This observation did not hold true for slightly more distant genes (<1kb), suggesting that tRNA expression only affects the most proximal genes. The removal of introns from genes in the synthetic genome also did not appear to greatly affect their expression levels (Figure S28). Overall, the design features of the syn6.5 genome appear to have only modest effects on transcript isoform boundaries and expression levels, making this strain a useful background to examine the effects of future SCRaMbLE perturbations.

### Morphology of yeast cells with syn6.5

To evaluate the cell morphology of yeast strains with multiple synthetic chromosomes, we visualized dividing syn6.5 cells with scanning electron microscopy (Figure 5I and J). They showed active multiplication and normal cell morphology, length and shape. The budding of daughter cells left ring-shaped bug scars on the cell wall of mother cells. We observed several cells at various stage of budding, including an aged mother cell with seven bud scars that was still actively budding. Rewriting multiple chromosomes as synthetic does not appear to affect cell morphology or cellular lifetime.

### Transferring *synIV* into synthetic chromosome strains using chromosome swapping

Currently, all yeast chromosomes have been synthesized separately in their individual host strains. In order to consolidate them more efficiently in the current syn6.5 strain, we deployed a new consolidation strategy: chromosome swapping (McCulloch et al., 2022). We hoped to develop a method to directly transfer individual new chromosomes to a recipient haploid strain that already carries multiple synthetic chromosomes. In yeast, *kar1-1* or *kar1-Δ15* mutations prevent nuclear fusion during mating when it is present in either parent (Conde and Fink, 1976). Thus, most progeny of these crosses remain haploid, but have a mixed cytoplasm. In these cells, chromosomes are occasionally transferred from the donor to the recipient strain. This process, called “exceptional cytoduction” (Dutcher, 1981) or chromoduction (Ji et al., 1993), results in an n+1 cell which can be selected for using proper auxotrophic and drug resistance markers.

Based on this phenomenon, a chromosome swapping method was developed entailing two steps: 1) introduction of the chromosome of interest into a recipient strain by chromoduction, resulting in an n+1 strain, and 2) destabilization of the native chromosome by inducing transcription through its centromere in the n+1 strain (McCulloch et al., 2022). To demonstrate how chromosome swapping could be deployed in synthetic chromosome consolidation, we picked the largest synthetic chromosome, *synIV*, as a “worst case scenario” proof of principle (Figure 6). The efficiency of chromoduction is inversely correlated with the chromosome size, and using this approach, each subsequent smaller synthetic chromosome swap is predicted to be even more efficient (Dutcher, 1981).

**Figure 6.**
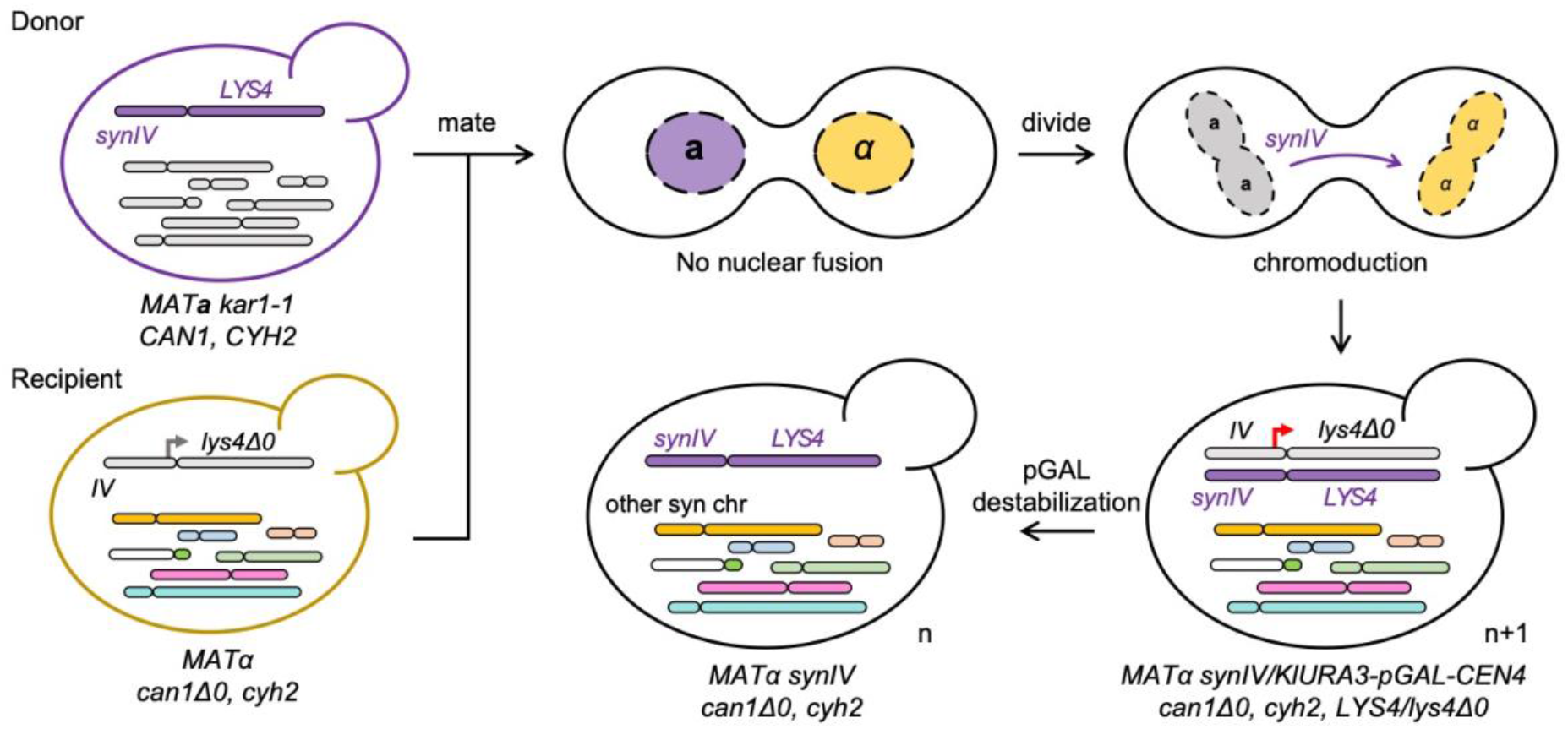
Consolidation of *synIV* into syn6.5 strains using chromosome swap. Donor: a strain carrying a synthetic chromosome(s) to be introduced with a selectable marker (*synIV* and *LYS4* in this example). Recipient: the strain of the opposite mating type already containing one or more multiple synthetic chromosomes, but retaining the native counterpart of the target (native *chrIV* and *lys4Δ0* here), which is tagged with *KlURA3* and pGAL-CEN (black hooked arrow). After cell conjugation, nuclear fusion is blocked by the *kar1-1* mutation. Transfer of *synIV* by chromoduction into recipient haploid progeny can be selected using the appropriate auxotrophic and drug-resistance markers. Finally, native *chrIV* is destabilized by induction of the pGAL promoter (red hooked arrow), which can be selected as 5-FOA^R^ due to the loss of *KlURA3*, completing the process of chromosome swapping.

First, in the recipient strain (syn6.5 as described above), we sequentially introduced 1) a *can1Δ0* deletion, 2) a *cyh2* mutation and 3) a *lys4Δ0* deletion in *chrIV* using CRISPR/Cas9, making the strain resistant to canavanine and cycloheximide and auxotrophic for lysine, respectively (Strain YZY402). The *can1Δ0* and *cyh2* markers are used to select against non-mating donor strains and occasionally formed diploid zygotes. The *lys4Δ0* can be complemented by *LYS4*^*+*^ in *synIV* from the donor strain. Next, the *kar1-1* mutation was introduced using CRISPR/Cas12a. After mating and selection for chromoductants, haploid progeny with both native and synthetic chromosome *IV* were successfully obtained. Finally, native *chrIV* was destabilized upon induction of the pGAL promoter and selection for 5-FOA^R^.

This transfer produced a yeast strain with more than half of its genome synthetic (Figure 6). This new strain (*synII, synIII, synIV, synV, synVI, synIXR, synX, synXII*) was confirmed as a haploid synthetic strain by WGS and flow cytometry (Figure S29). It was characterized by slower growth, with a G1 arrest, suggesting a cell cycle defect, and will require further bug mapping to generate a high fitness derivative. Combined with CRISPR D-BUGS, chromosome swapping is a NextGen strategy to consolidate and debug new, incoming synthetic chromosomes into the genetic background of other synthetic ones.

## Discussions

The goal of Sc2.0 project is to build the first eukaryotic organism with a completely redesigned and human-synthesized genome. One yeast strain carrying 6.5 synthetic chromosomes was successfully constructed from several rounds of endoreduplication intercross, followed by extensive debugging. We also successfully used chromosome swapping to incorporate *synIV* creating the syn7.5 multiple synthetic strain. Continued use of these methods will facilitate consolidation of all remaining synthetic chromosomes to construct a final, fully synthetic Sc2.0 strain.

In Sc2.0, thousands of genome-wide edits were introduced as designer features, including removal of introns and mobile elements, stop codon swaps intended to remove all instances of the TAG stop codon, addition of loxPsym sites, and relocation of all tRNA genes from their native loci onto a neochromosome. These features facilitate a variety of applications such as genome minimization, novel amino acid incorporation and SCRaMbLE. As an inducible evolution system, SCRaMbLE has been used to exploit genomic structural variation, biosynthesis pathway engineering and host strain improvement (Blount et al., 2018; Luo et al., 2018b; Shen et al., 2018; Shen et al., 2016; Zhao et al., 2020). Most recently, combined with deep transcript isoform profiling, synthetic chromosomes and SCRaMbLE have been used to study genome architecture and its contribution to transcriptional regulation (Brooks *et al*., 2022). Genome-wide SCRaMbLE and rearrangements also provide insights into 3D spatial organization and chromatin-accessibility (Zhou et al., 2022). With more synthetic chromosomes incorporated, SCRaMbLE can also be deployed to study additional biological questions such as new yeast phenotypes and minimized chromosomes and genomes, making Sc2.0 a novel platform to understand eukaryotic genomes and develop industrial applications.

Native *S. cerevisiae* contains 295 introns that belong to 280 genes, and 91 of these introns were deleted in the syn6.5 strains. Previous studies have demonstrated that intron accumulation might aid yeast starvation response by sequestering available splicing factors and affecting splicing of RNAs encoded by other genes, especially ribosomal-protein genes, which are hypothesized to be regulated by rapid intron removal once fresh nutrients are supplied (Morgan et al., 2019; Parenteau et al., 2019). As the splicing machinery is presumably more readily available in the syn6.5 strain, it would be interesting to evaluate the impact its reduced intron content may have on splice isoforms and spliced/unspliced ratios of the genes residing on the residual native chromosomes in the future.

The introduction of these designer edits also results in unexpected bugs in the form of a wide variety of growth defects. We developed a new mapping strategy CRISPR D-BUGS to map these bugs, facilitating their elimination by reversion. Interestingly, several bugs found in this and companion studies map to loxPsym sites introduced downstream of what are now classified as “dubious ORFs” which end up damaging promoters and 5’ UTRs of authentic genes. Thus, this is a type of bug that results from the rules of Open Reading Frame annotations which were adopted early in the Sc2.0 project, when dubious ORFs were not yet well defined and annotated.

This debugging method will also be helpful during further consolidation of additional synthetic chromosomes. Using CRISPR-D-BUGs, we first mapped a *synII* growth defect to two repetitive loxPsym sites in the *SHM1* 3’ UTR, which led to reduced expression by an unknown mechanism. *SHM1* encodes mitochondrial serine hydroxymethyltransferase (SHMT), an enzyme that can interconvert serine and glycine and produce 5,10 methylene tetrahydrofolates (CH2-THF). Shm1p only comprises about 5% of total SHMT activity whereas its cytosolic isoform Shm2p comprises the majority (Kastanos et al., 1997). Shm1p and Shm2p are conserved from bacteria to humans, and both enzymes are essential components in the one-carbon metabolism cycle, which provides a crucial substrate for mitochondria initiator tRNA formylation and other reactions required for biosynthesis of nucleotides, amino acids and lipids (Lee et al., 2013; Piper et al., 2000). The *shm1Δ* strain was shown to have a near-WT level of formylation and mitochondrial protein expression, suggesting that cytosolic SHMT activity produces a sufficient supply of one carbon units in the absence of Shm1p (May *et al*., 2020).

A further clue about this bug came from the observation that an extra copy of *TSC10* suppressed the *synII* growth phenotype. We found that overexpressing *TSC10* either from a plasmid or via ectopic integration at the *HO* locus rescued the growth defect of *synII* (Figure S30). *TSC10* is an essential gene, encoding a 3-ketosphinganine reductase catalyzing the second step in phytosphingosine synthesis, using serine as the key precursor molecule (Beeler et al., 1998). This pathway is the basis for all sphingolipids made in yeast, including ceramide, a major component of the mitochondrial membrane. Interestingly, this pathway is highly involved in heat stress response and activated immediately upon onset of heat stress (Chen et al., 2015). Several sphingolipids were also implicated as secondary messengers involved in signaling pathways that regulate the heat stress response (Jenkins et al., 1997). Based on this, we speculate that in the original *synII* strain (chr02_9_03), sphingolipid/ceramide biosynthesis was also affected when grown on YPG medium due to a deficit of cytosolic serine. This is supported by the fact that serine is mainly synthesized from glycine via SHMT activity of Shm1p and Shm2p in a “gluconeogenic” pathway on non-fermentable glycerol but not on glucose (Albers et al., 2003; Melcher and Entian, 1992). In short, we hypothesize that *SHM1* under-expression leads to insufficient ceramide biosynthesis and at high temperature this reduces mitochondrial function.

During consolidation, we noted the existence of a combinatorial bug revealed when combining *synIII* and *synX*, which was later mapped and resolved using CRISPR D-BUGS. Individual *synIII* or *synX* strains are highly fit. Using a *synIII, synX* strain, we successfully mapped the combinatorial bug to *synSUP61*, which under-expresses tRNA_Ser_^CGA^ and *SWI3*, which is under-expressed due to a loxPsym site in its 5’ UTR, respectively. Reduction of *SWI3* expression and consequently, inositol auxotrophy phenotypes were further enhanced by reduced tRNA_Ser_^CGA^ abundance. In support of this model, mutation of tandem codons recognized by tRNA_Ser_^CGA^ in *SWI3* dramatically suppressed the phenotype. tRNA abundance and codon usage are closely linked, with “rich” codons corresponding to abundant tRNAs overrepresented in highly expressed genes (Frumkin et al., 2018; Wei et al., 2019). Optimal codon usage is also predicted to ensure the proper speed of translation elongation for efficiency, accuracy and correct protein folding (Guimaraes *et al*., 2020; Liu, 2020). Under stressful conditions, yeast can alter tRNA abundance to facilitate selective stress-related translation and interestingly, rare codons corresponding to low-abundance tRNAs are enriched in stress-responsive genes potentially allowing for efficient and sensitive regulation (Torrent et al., 2018). From a synthetic chromosome consolidation perspective, we discovered a completely unexpected connection between abundance of a single copy tRNA, inositol auxotroph and potentially, chromatin dynamics, consistent with regulation via codon usage and tRNA pool adjustments. In the current stage of Sc2.0, the tRNAs have been relocated into chromosome-specific tRNA arrays but will eventually be consolidated into a single neochromosome. Each tRNA gene in the arrays is flanked by rox recombination sites that can be recognized by Dre recombinase. This site-specific recombinase system can be deployed to perform a tRNA gene-specific form of SCRaMbLE, but it can also be deployed in the future to simultaneously remove all of the chromosome specific tRNA arrays once a future version of the tRNA neochromosome, namely one entirely lacking rox sites, is introduced into the Sc2.0 progenitor strain.

Finally, we have constructed 21 strains with all pairwise combinations of 7 previously completed synthetic chromosomes (Figure S31). Happily, the vast majority of these show no growth defect at 30°C. However, we noted modest growth defects at 37° suggesting that additional but mild combinatorial bugs may exist in the strains containing *synII/synXII* and *synII/synVI*. Consequently, the current syn6.5 strain containing these synthetic chromosomes still shows a growth defect at high temperature, although the fitness has improved dramatically since fixing these known bugs. Debugging combinatorial defects is more challenging than single bugs, but we are confident that CRISPR D-BUGS will greatly facilitate deciphering these.

## Experimental Procedures

### Synthetic chromosome versions

We started the consolidation using strains containing individual synthetic chromosome and when necessary, we switched the mating type using CRISPR (Xie et al., 2018). Detailed information on intermediate strain names, genotypes and version numbers are listed in Table S4. Briefly, they are *synII* yeast_chr02_9_03 (Shen *et al*., 2017), *synIII* yeast_chr03_9_02 (Annaluru *et al*., 2014), synIV yeast_chr04_9_03 (Zhang *et al*., 2022), *synV* yeast_chr05_9_22 (Xie *et al*., 2017), *synVI* yeast_chr06_9_03 (Mitchell *et al*., 2017), *synIXR* genebank JN020955 (Dymond *et al*., 2011), *synX* yeast_chr10_9_01 (Wu *et al*., 2017) and *synXII* yeast_chr12_9_02 (Zhang *et al*., 2017).

### Yeast media, growth and transformation

Yeast strains were cultured in YPD-rich medium or defined SC media with appropriate components dropped out. All yeast transformations were performed using standard lithium acetate protocols (Brachmann et al., 1998). To check *synII* growth defects, YPG plates contained 3% glycerol. For inositol auxotrophy tests, the inositol free YNB medium was prepared using yeast nitrogen base w/o inositol (US Biological Y2030-01), and supplemented with 5 g/L ammonium sulfate, 2% dextrose and the necessary amino acid supplements. Control plates were supplemented with 76 mg/L myo-inositol (Sigma I5125).

### Consolidation using endoreduplication intercross

Two separate synthetic chromosomes were consolidated using endoreduplication intercross from their host strain with opposite mating types (Mitchell *et al*., 2017). First, the two host haploid strains were engineered: the *KlURA3*-pGAL-CENx module was integrated into the native counterparts of the target synthetic chromosomes to be lost (Hill and Bloom, 1987). Our early experiments suggested that destabilizing two native chromosomes per diploid by this method worked more reliably than trying to destabilize larger numbers of native chromosomes at the same time. After mating and inoculation in YP+Galactose (2%) medium overnight, the relevant heterozygous diploid strain was screened on SC+5-FOA plates for successful destabilization of both target chromosomes, generating a 2n-2 strain. After growth for 24 hours allowing for endoreduplication in YPD, the strain was cultured in sporulation medium at room temperature. Finally, from tetrad dissection, spore clones with more than both target synthetic chromosomes were obtained. This process was continued to consolidate more synthetic chromosomes (Figure S1).

### tRNA array design and integration

For each synthetic chromosome, a tRNA array containing all the synthetic version of tRNA genes from its host chromosome was constructed (Table S2). Each syn-tRNA contains 500bp 5’ and 40bp 3’ flanking sequences from *Eremothecium* (*Ashbya*) *gossypii* or *Eremothecium cymbalariae*. The tRNA arrays were integrated was described in Figure S2. The arrays were released from plasmids using restriction enzyme digestion (Table S3). To integrate tRNA arrays, we constructed junction DNAs with 500bp homology arms to the target genomic locus, 500bp homology arms to the linearized tRNA array and the *KlURA3* selection marker at one end. Integrations were selected on SC–Ura plates and confirmed by colony PCR. Afterwards, the *KlURA3* marker was deleted using CRISPR/Cas9 and a gRNA.KlURA3 (ACCAGTAACCCCGTGGGCGT), provided with a flanking donor DNA.

### CRISPR D-BUGS

In this study, we developed CRISPR D-BUGS to quickly and reliably map the bug on a synthetic chromosome. Step one in this process is to determine whether the fitness defect is recessive (most cases) or dominant. Assuming that the defect to be mapped is recessive, we first created a diploid strain of yeast heterozygous for the target chromosome arm with a *URA3* marker integrated in an intergenic region close to the telomere (∼25kb) of the native chromosome. Then several gRNAs targeting WT PCRtags at different regions along the chromosome were selected (Table S5). For the initial round of CRISPR D-BUGS, it is good to have gRNAs targeting near the telomeres, near the middle of the left or right arms, and on either side of the centromere (at least ∼10kb away). The gRNA was assembled into a CRISPR/pGAL-Cas9 plasmid backbone (pYZ555 with *LEU2* marker) using Golden Gate cloning (Zhao and Boeke, 2020).

Afterwards, the heterozygous diploid strain was transformed with the CRISPR plasmid and selected on SC–Leu dextrose plates (Cas9 OFF). A single colony was inoculated in SC–Leu galactose medium and incubated at 30°C overnight (Cas9 ON). The medium was diluted and plated on 5-FOA plates to select the single colonies with successful mitotic recombination, further confirmed using PCRtag assays. Finally, we assessed fitness by single colony formation spot tests to identify the fitness boundary. Once the fitness boundary is rough-mapped, further intermediate gRNAs can be chosen for fine mapping until a gRNA that produces a mix of fit and unfit clones is identified. WGS of the fit and unfit clones can then be deployed to fine-map the fitness boundary.

### Genomic editing using CRISPR/SpCas9-NG

The CRISPR/SpCas9-NG system was used to repair an accidental single base pair mutation in *synI YAL061W*. The SpCas9-NG ORF was subcloned from pX330-SpCas9-NG obtained from Addgene (#117919), and assembled with *TEF1* promoter and *CYC1* terminator (Zhang et al., 2022). Briefly, the gRNA (GGTCCATGTGCTACACACAC) targeting at *YAL061W* with CG as the PAM was used to repair the mutation in yJL663 with a draft version of *synI*. The donor DNA was the PCR product from wild-type genomic DNA containing 140bp homology arm on each side of the target mutation. We got 3 out of 11 positive colonies where the mutation was repaired.

### Genomic editing using CRISPR/Cas9 and Cas12a

We also used CRISPR/Cas9 and Cas12a (also called Cpf1) to repair the mapped bugs or introduce new variants. We followed the protocols as described previously (DiCarlo *et al*., 2013; Swiat et al., 2017). All targets, gRNA and PAM sequences for this study are listed in Table S6.

### Pulsed-field gel electrophoresis (PFGE)

To evaluate the multiple synthetic chromosomes in syn6.5 strains, chromosome plugs were prepared the separated by clamped homogeneous electric field (CHEF) gel electrophoresis using the CHEF-DR III Pulsed-Field Electrophoresis System (Bio-Rad), as previously described (Luo et al., 2018a). The following program was used, temperature: 14 °C, voltage: 6 V/cm, switch time: 60 s to 120 s, run time: 20 h, included angle: 120°, using 0.5 × Tris-Borate-EDTA buffer and a 1% gel with low melting point agarose (Bio-Rad #1620137). Gels were stained with 5 μg/ml ethidium bromide in water after electrophoresis for 30 min, de-stained in water for 30 min, and then imaged.

### Whole genome sequencing and alignment

The yeast genomic DNA samples for sequencing were prepared using a Norgen Biotek fungi/yeast genomic DNA isolation kit (Cat No. 27300). The library was prepared using NEBNext Ultra II FS DNA library prep kit (NEB E7805L) with 500 ng genomic DNA as input. The whole genome sequencing was performed using an Illumina 4000 system using pair-end 36bp protocol. All raw reads were trimmed to remove adaptor sequence using Trimmomatic, and subsequently mapped to synthetic chromosome sequences using bowtie2 software. The coverage for each locus was calculated using BedTools and normalized to average genome-wide coverage.

### GFP tagging and immunoblotting

To quantify protein expression level of *SHM1* by immunoblotting, we first tagged it with GFP. Since the loxPsym sites are inserted in 3’ UTR close to the stop codon, we integrated the tag at N-terminal instead of C-terminal. We used the same sequence and design of SWAT library (Weill et al., 2018). We first isolated the strain of *SHM1* tagged with GFP at N terminal and its native promoter from the SWAT library. Then, we PCR amplified the region containing GFP and 500bp homology arms on each side from its genomic DNA as the donor DNA, which was transformed with CRISPR/Cas9, using one gRNA (GACTAGCGATTGTGCACCAC). Successful integration was confirmed with colony PCR and Sanger sequencing. Notably, the mitochondria targeting signal of Shm1p was not affected.

We tagged *SHM1* with GFP in original *synII* (9.03) and fixed *synII* (9.04) strain, generating YZY516 and YZY517, respectively. The original wild-type strain from SWAT library, YZY208, was used as a control. These strains were cultured in YPD medium overnight, and then diluted to produce a log phase culture. The cell lysate was prepared and run on SDS-PAGE as previously described (Ikushima et al., 2015). Proteins were transferred to a PVDF membrane for blocking, antibody binding and imaging. Anti-GFP antibody from mouse (Roche 11814460001) at 1:1000 dilution and anti-H3 antibody from rabbit (Abcam ab1791) at 1:2500 dilution were used as the primary antibodies. IRDye Goat anti-mouse IgG (LI-COR Biosciences 926-32210) and Goat anti-rabbit IgG (LI-COR Biosciences 926-68071) were used as the secondary antibodies, respectively. The fluorescence signal was detected on an Odyssey CLx Imager from LI-COR.

To quantify protein expression level of *SWI3* with loxPsym in 5’ UTR and/or 3’ UTR at synX, we first tagged it with 3×Flag tags (DYKDHDGDYKDHDIDYKDDDDK) and a GS linker (GGGGS)_3_ at C-terminus. The immunoblotting was performed with the same method as above. Anti-Flag antibody from mouse (Sigma F1804) at 1:2000 dilution was used as the primary antibodies. The same internal control and secondary antibodies were used as above.

### Real-time PCR

We used real-time PCR to check the expression level of *SHM1* as previously described (Mitchell *et al*., 2017). Briefly, from 3 single colonies as triplicates, the RNA was prepared using RNeasy Mini kit (Qiagen 74106). First strand cDNA was prepared using SuperScript IV Reverse Transcriptase (Invitrogen 18090050) and oligo d(T)_20_ primer. The expression level was tested using Lightcycler 480 SYBR Green I Master Mix (Lightcycler 04887352001) in a 10 μl reaction system. The qPCR was performed and analyzed using the LightCycler 480 System. For *SHM1*, the forward primer (GCTCTGGAACTGTACGGATTA) and reverse primer (ACGTTCATGATAGCGGAGTAAA) were designed by IDT PrimerQuest Tool. The *TAF10* was used as the internal control (Teste et al., 2009).

### RNA extraction for transcript profiling

Total RNA was extracted from 50mL flash-frozen cell pellets grown to mid-log (OD_600_ ∼0.65-0.85) using MasterPure™ yeast RNA purification kit (Lucigen) including a DNaseI treatment step. RNA (diluted 1:10) quality and concentration were measured by Agilent 2100 Bioanalyzer with the Agilent RNA 6000 Nano Kit and Qubit™ RNA High Sensitivity Kit (Thermo Fisher), respectively.

### Direct RNA sequencing

Poly(A) mRNA was enriched from 93.75 µg total RNA on 250 µL Dynabeads oligo(dT)_25_ beads. The direct RNA sequencing kit (SQK-RNA002, Oxford Nanopore Technology) was used to generate libraries from 500 ng poly(A) RNA. An optional reverse transcription was performed at 50°C for 50 min using SuperScript™ IV reverse transcriptase (Invitrogen) in between the ligation of the RTA and RMX adaptors. Following reverse transcription the RNA:cDNA was cleaned up with 1.8 volumes of Agencourt RNAclean XP beads and washed with 70% ethanol. Following RMX ligation only 1 volume of beads were used in the clean-up, and WSB (SQK-RNA002) was used in the wash steps. Direct RNA libraries (typically 150-200 ng) were loaded onto primed (EXP-FLP001) MinION flow cells (FLO-MIN106D, R9 version) in RRB buffer and run on the GridION with MinKNOW 3.1.8 for up to 72 hours.

### Directional mRNA sequencing

NEBNext® Poly(A) mRNA magnetic isolation module (E7490) was used to enrich poly(A) mRNA from 500 ng total RNA with 5 µL 1:500 diluted ERCC RNA Spike-In control mix (Thermo Fisher) in 50 µL. The NEBNext® Ultra™ II directional RNA library prep kit for Illumina with sample purification beads (E7765) was used to prepare stranded mRNA sequencing libraries from the poly(A) RNA. Libraries were amplified for 11 cycles with i7 index primers (E7500S). Libraries were individually cleaned-up with 0.9 volumes of sample purification beads and concentration and size distributions were measured by Qubit™ dsDNA high sensitivity kit and by Agilent 2100 Bioanalyzer with the Agilent high sensitivity DNA kit. Equimolar amounts were combined of all samples, cleaned-up on 0.9 volumes of sample purification beads, and submitted for 150bp paired end sequencing on the NextSeq 500 (Illumina) at the EMBL Genomics Core.

### Base calling, quality-filtering, and long-read alignment

Nanopore long reads were base-called, trimmed of adapter sequences, and filtered for quality, retaining only those with the best alignment scores for multi-mapping reads, as previously described (Brooks *et al*., 2022).

### Gene expression quantification

NEBNext UltraII directional mRNA was quantified at the transcript-level by Salmon v1.6.0 (Patro et al., 2017), aligning to known transcripts as well as those identified in the long-read sequencing. When observing mature mRNA transcripts, reads were aligned against a database of transcripts with and without introns. Salmon was run with sequence and position bias modelling enabled. Differential gene expression analysis was performed in DESeq2 (Love et al., 2014).

### Chromosome swapping to consolidate *synIV*

We developed a method to directly consolidate individual new chromosome with other multiple synthetic chromosomes. In the recipient strain with 6.5 synthetic chromosomes, *can1Δ0, lys4Δ0* with ORF deleted, *cyh2* mutation (Q38K) were introduced using stepwise CRISPR/Cas9 editing with donor DNA provided. Native *chrIV* was targeted with the *KlURA3-pGAL-CEN4* module. The *kar1-1* mutation (P150S) was introduced using CRISPR/Cas12a, generating the final recipient strain YZY402.

The donor strain (yWZ675, carrying *synIV*, yeast_chr04_9_03) and recipient strain (YZY402, carrying 6.5 synthetic chromosomes) were prepared as fresh patches (∼2 cm in diameter) on separate YPD plates incubated overnight at 30°C, and then mated together by replica plating.

After incubation at 30°C for 12 h, the mating plate was replica-plated to a selection plate of SC– Lys+Can (60 ng/µl) +cycloheximide (10 ng/µl). After incubation at 30°C for one week, haploid progeny with both *chrIV* and *synIV* (n+1) were successfully obtained and re-streaked to new selection plates, which were then checked by PCRtag assays. The efficiency for *synIV* transfer was around 10% (2 out of 23 screened). Finally, the strain was incubated in YP+galactose medium to destabilize the native chromosome and selected on 5-FOA plates, generating the final haploid strain consolidated with new synthetic chromosomes.

### DNA content assay

We used a previously described DNA content assay (Haase and Lew, 1997). Briefly, about 5 × 10^6^ cells were fixed in ethanol 70% for 1 h at room temperature, then pelleted, washed, and incubated in 10 mM Tris pH 7.5 with RNase A (0.1 mg/ml, ThermoFisher EN0531) for 2 h at 37°C. Cells were pelleted, resuspended in 10 mM Tris pH7.5 with propidium iodide (5 µg/ml, Invitrogen P3566), and incubated for 1 h in dark at 4 ºC. Finally, the cells were pelleted and resuspended in 0.5 ml 50 mM Tris pH 7.5, and analyzed using a BD AccuriTM C6 flow cytometer.

### Scanning electron microscopy

Cultured yeast cells were plated on 12mm poly-l-lysine coated glass coverslip in 24 well dish and fixed with 2.5% glutaraldehyde in PBS for one hour. After washing with PBS, the yeast cells were post fixed in 1% Osmium tetroxide for one hour, dehydrated in a series of ethanol solutions (30%, 50%, 70%, 85%, 95%, 100%), and dried with Tousimis autosamdsri 931 (Rockwille, MD) critical point dryer. The coverslips were put on SEM stabs, sputter coated with gold/palladium by DESK V TSC HP Denton Vacuum (Moorestown, NJ), and imaged by Zeiss Gemini300 FESEM (Carl Zeiss Microscopy, Oberkochen, Germany) using secondary electron detector at 3 kv with working distance 10 mm.

## Acknowledgements

We thank M.J. Sadhu, Pan Cheng, Guangbin Shi and Michael Pacold for helpful discussions. We thank Megan Hogan, Raven Luther, Hannah Ashe and Gwen Ellis from the Matthew Maurano lab for assistance with WGS. We give special thanks to the Sc2.0 consortium for many discussions and collaborations. This work is part of Sc2.0 project (http://syntheticyeast.org/), supported by NSF grants MCB-1026068, MCB-1443299, MCB-1616111 and MCB-1921641 to JDB. Microscopy Laboratory is partially supported by Laura and Isaac Perlmutter Cancer Center Support Grant NIH/NCI P30CA016087, and Zeiss Gemini300SEM was purchased with NIH S10 OD019974. Transcriptome profiling was funded by a Volkswagen Stiftung grant (94769) to LMS.

