## Supplemental Figures for "Debugging and consolidating multiple synthetic chromosomes reveals combinatorial genetic interactions"

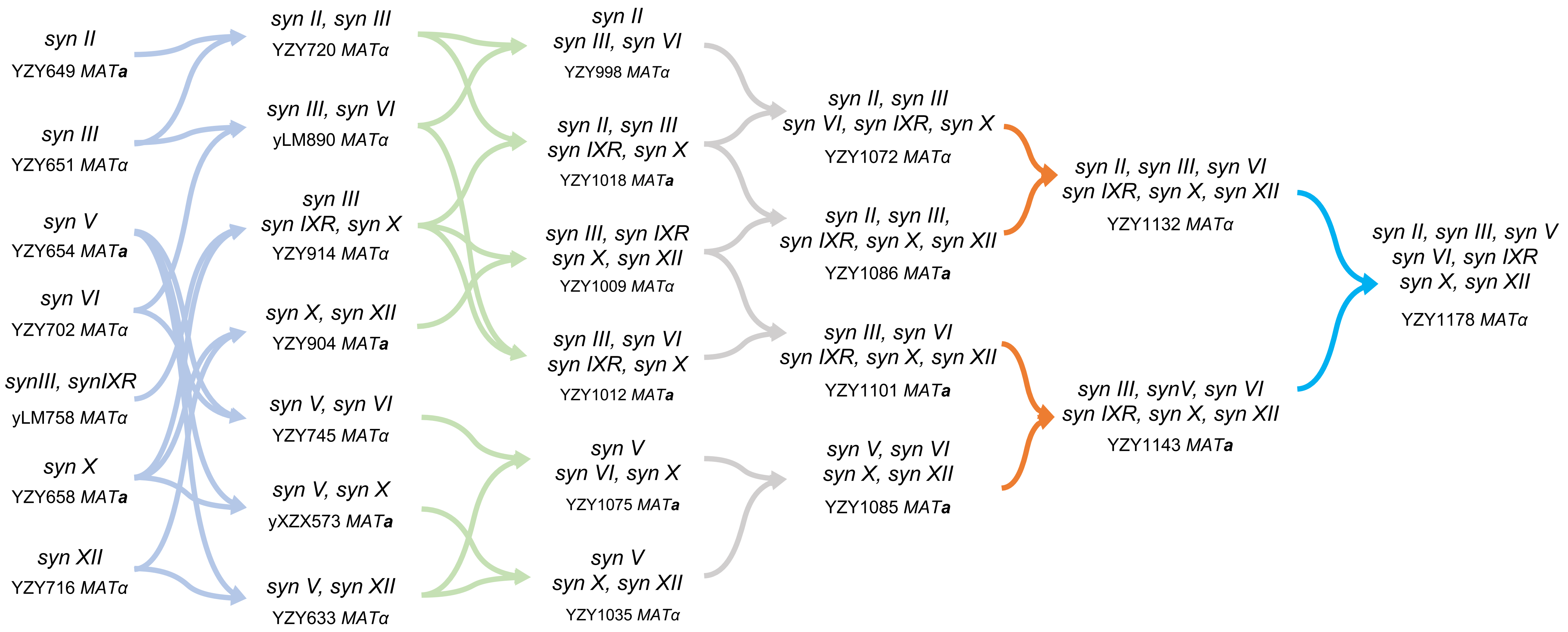


**Figure S1. The consolidation process using endoreduplication intercrossing.**

Two strains with different synthetic chromosomes and opposite mating type were mated together, generating heterozygous diploid strains.
See Table S2 for further details of the strains. As needed, mating types were switched using CRISPR (Xie et al., 2018).

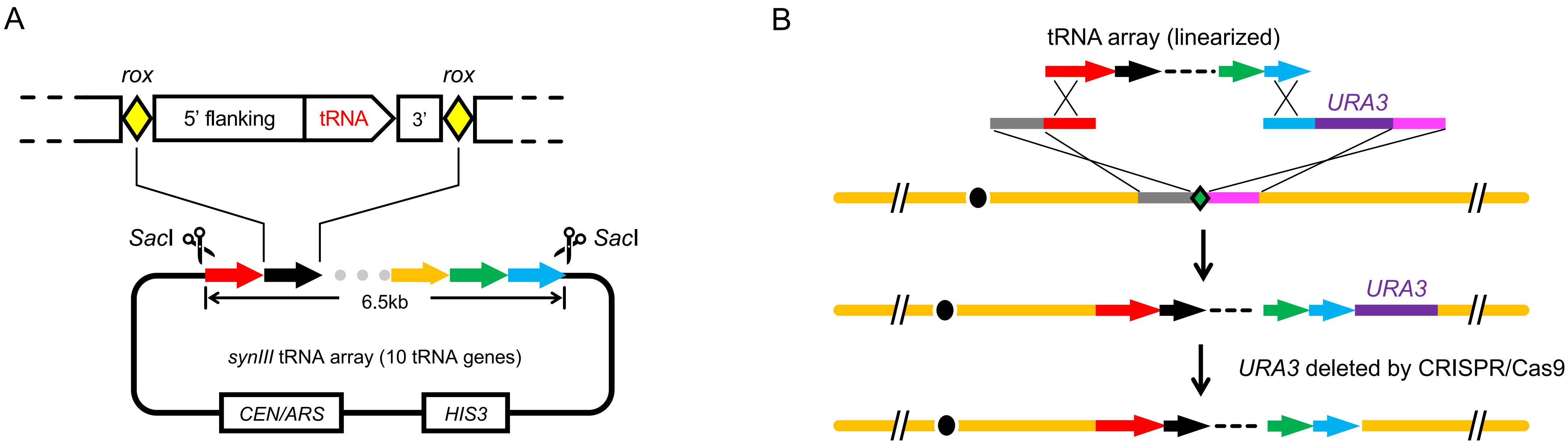


**Figure S2. Integration of tRNA arrays into each synthetic chromosome.**

(A) Each tRNA array contains all the tRNA genes from its native chromosome. The array is stored in a plasmid and can be released by restriction enzyme digestion. The *synIII* tRNA array is shown as an example here. The anatomy for each tRNA array is shown in Figure S3.

(B) The tRNA array was integrated into the synthetic chromosome using a two-step homologous recombination method. First, it was integrated and selected with *URA3* marker. Second, the *URA3* marker was deleted using CRISPR/Cas9 with donor provided. The restriction enzymes to linear tRNA array and integration loci are listed in Table S3.

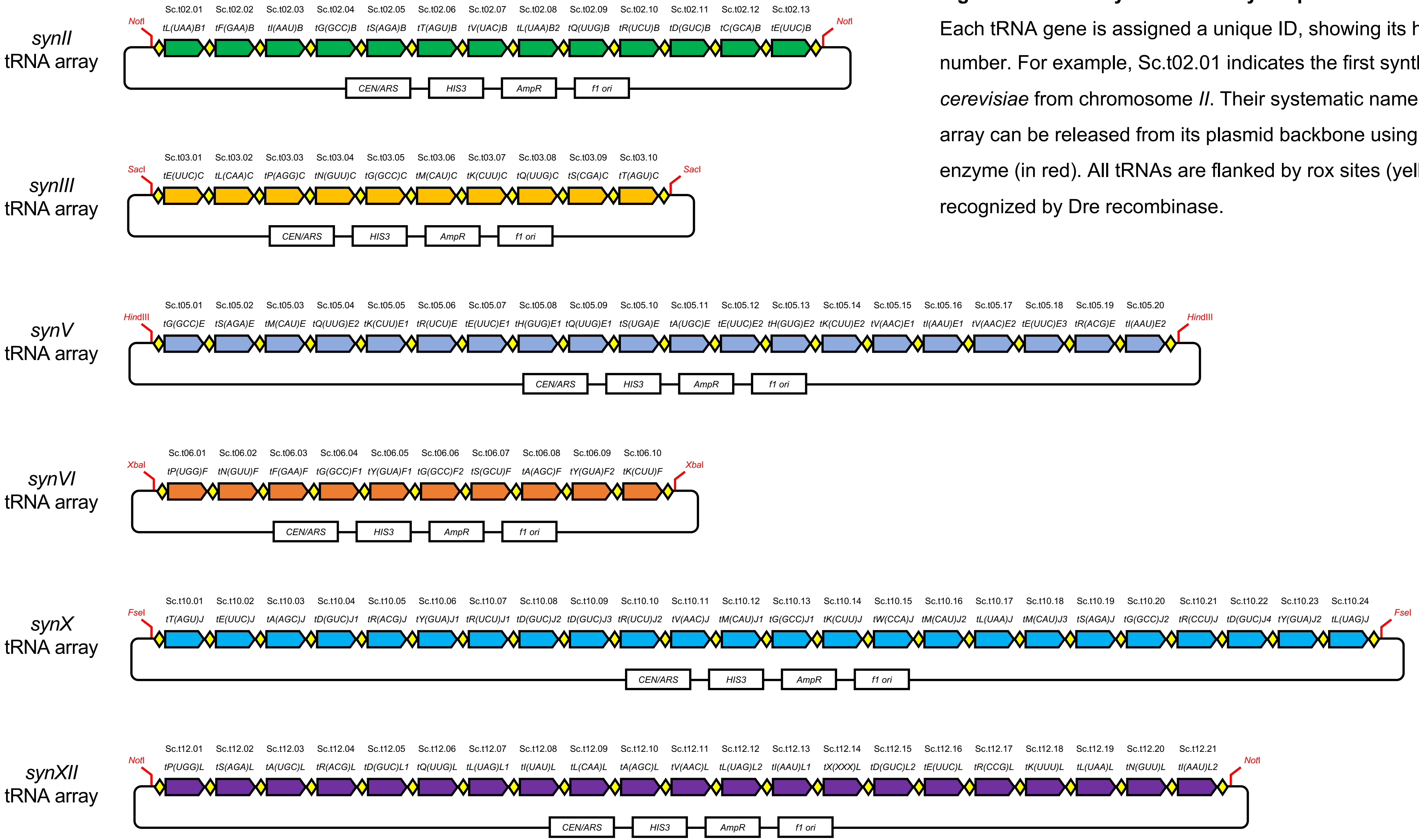


**Figure S3. Anatomy of tRNA arrays in plasmids.**

Each tRNA gene is assigned a unique ID, showing its host chromosome and gene number. For example, Sc.t02.01 indicates the first synthetic tRNA gene in *S. cerevisiae* from chromosome *II*. Their systematic names are also labelled. Each tRNA array can be released from its plasmid backbone using an appropriate restriction enzyme (in red). All tRNAs are flanked by rox sites (yellow diamond) which can be recognized by Dre recombinase.

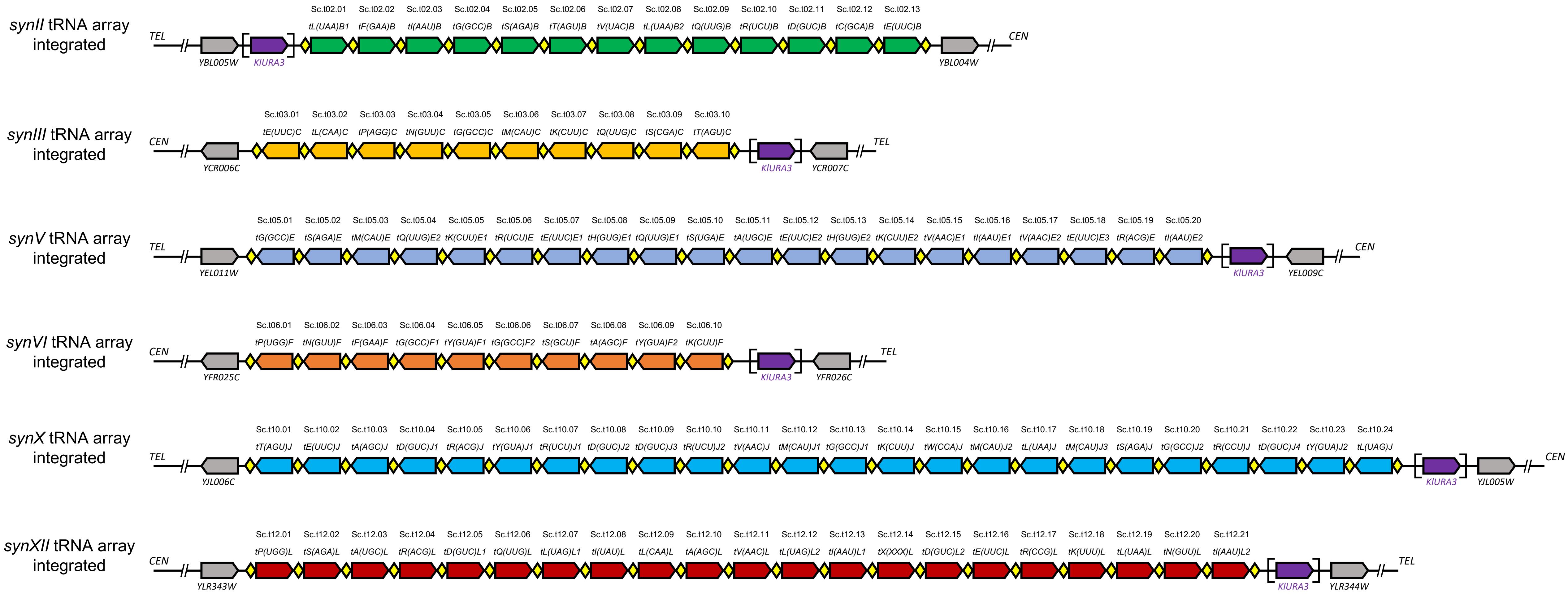


**Figure S4. Anatomy of tRNA arrays integrated in synthetic chromosomes.**

Each tRNA array was integrated into its host synthetic chromosomes, using the method in Figure S2. *KlURA3* (purple in square bracket) was used as an integration marker, which was then deleted by CRISPR/Cas9. The rox sites (yellow diamond) were also integrated, enabling tRNA array rearrangements in future applications.

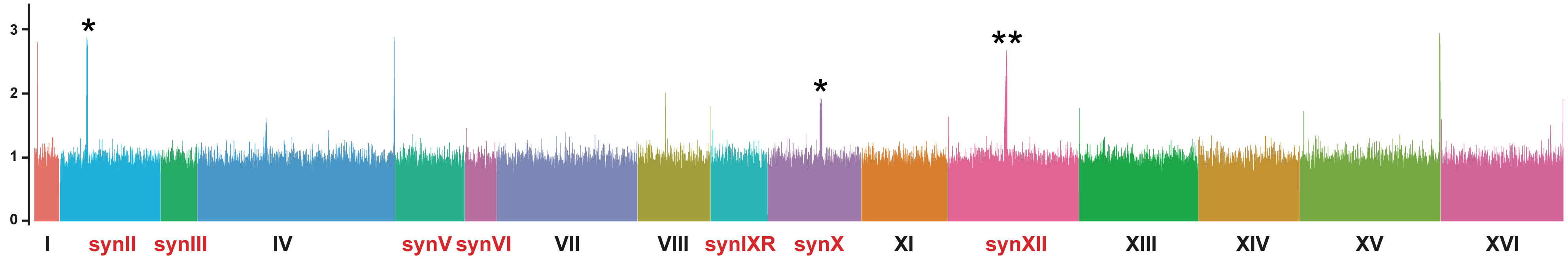


**Figure S5. Sequencing coverage in the WGS for multiple synthetic chromosomes.**

The presence of complete multiple synthetic chromosomes (*synII, synIII, synV, synVI, synIXR, synX, synXII*) were confirmed by whole genome sequencing (WGS).

* Duplications of tRNA happened during their initial integration, which were later repaired by CRISPR/Cas9.

** Higher coverage of rRNA repeats were detected in *synXII*.

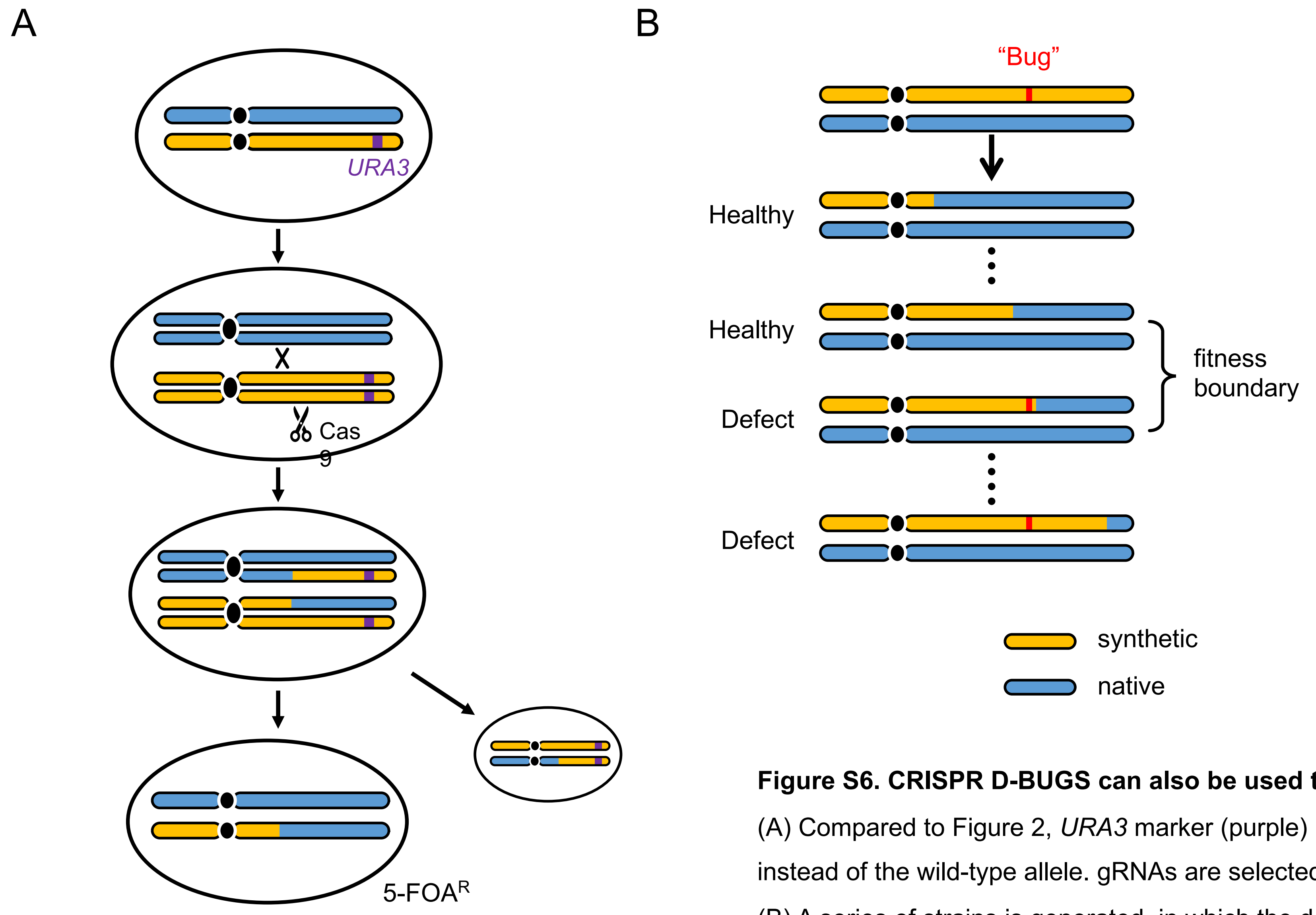


**Figure S6. CRISPR D-BUGS can also be used to map dominant bugs.**

(A) Compared to Figure 2, *URA3* marker (purple) is pre-integrated into the synthetic allele (orange), instead of the wild-type allele. gRNAs are selected to target Cas9 cleavage at synthetic PCRtags.

(B) A series of strains is generated, in which the defect phenotype indicates the presence of the dominant bug in the synthetic sequence.

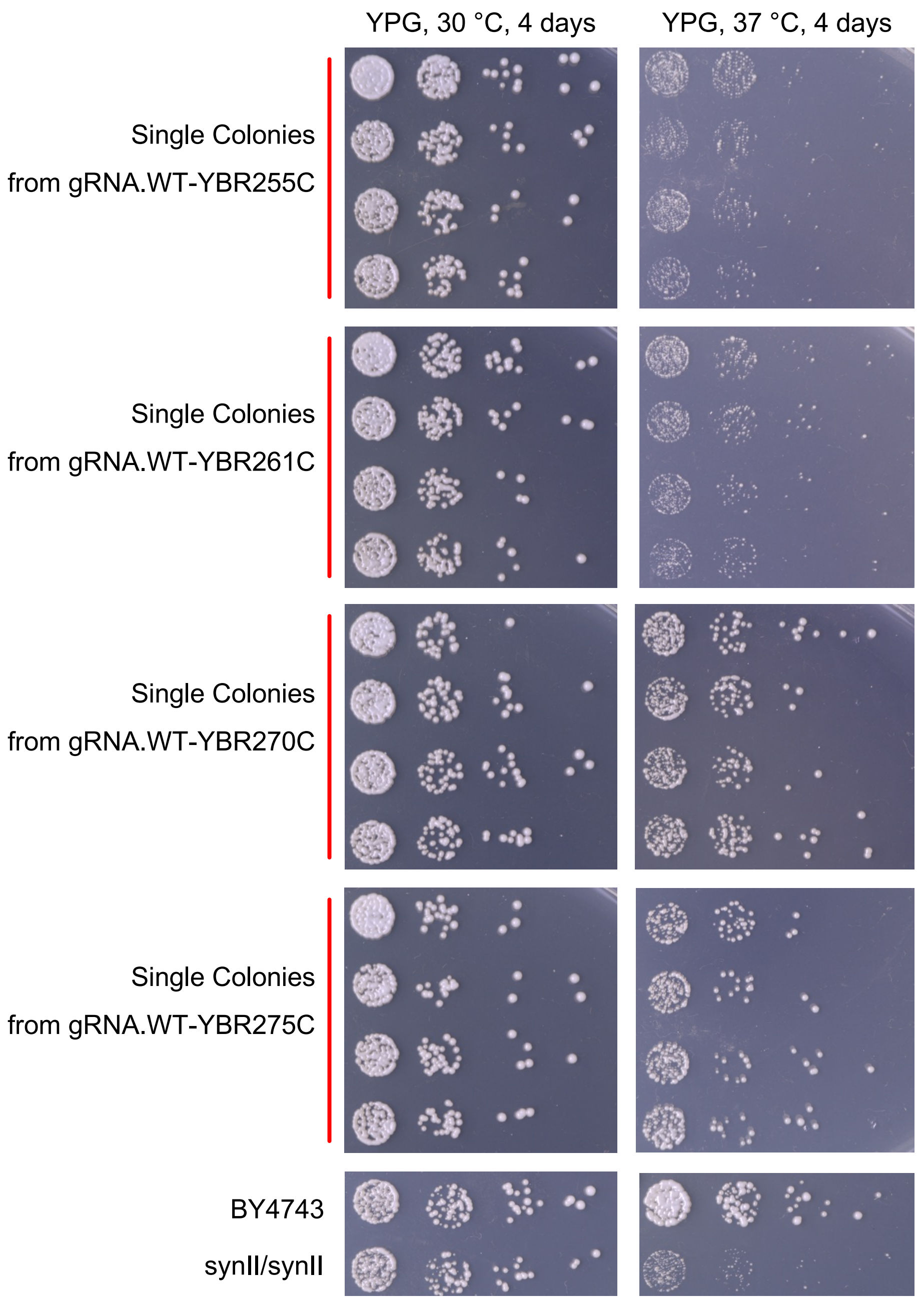


**Figure S7. Fitness of single colonies generated from the first round of *synII* bug mapping.**

The gRNA targets are labelled on the left side. For each group, all four single colonies showed consistent fitness level and only one was shown in Figure 3B.

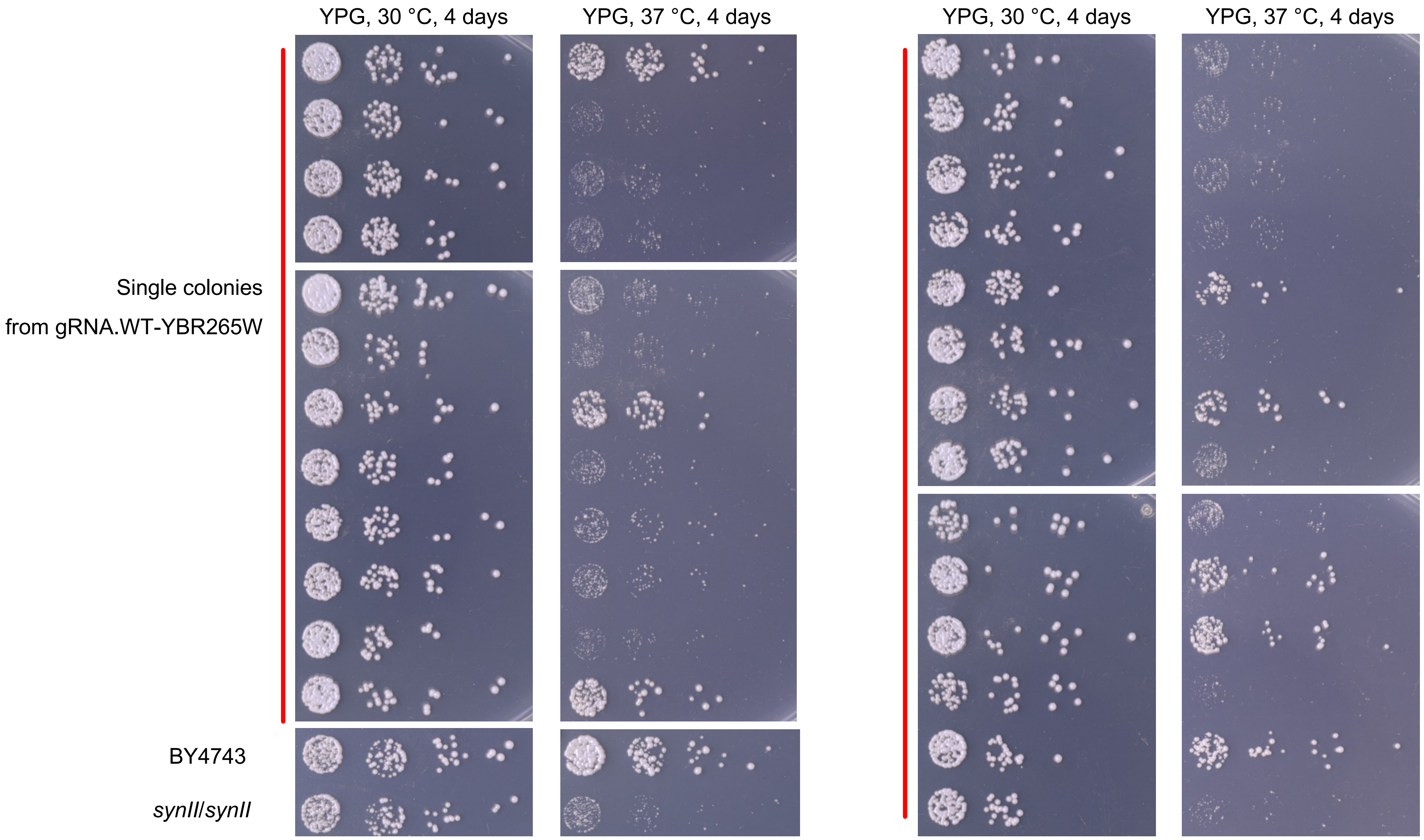


**Figure S8. Single colonies generated from the same gRNA.YBR265W showed a mixture of fitness levels.**

Single colonies were isolated from the same 5-FOA plate and showed variable growth phenotypes. Their recombination sites were mapped using WGS, aligning to synII as the reference, as in Figure 3C.

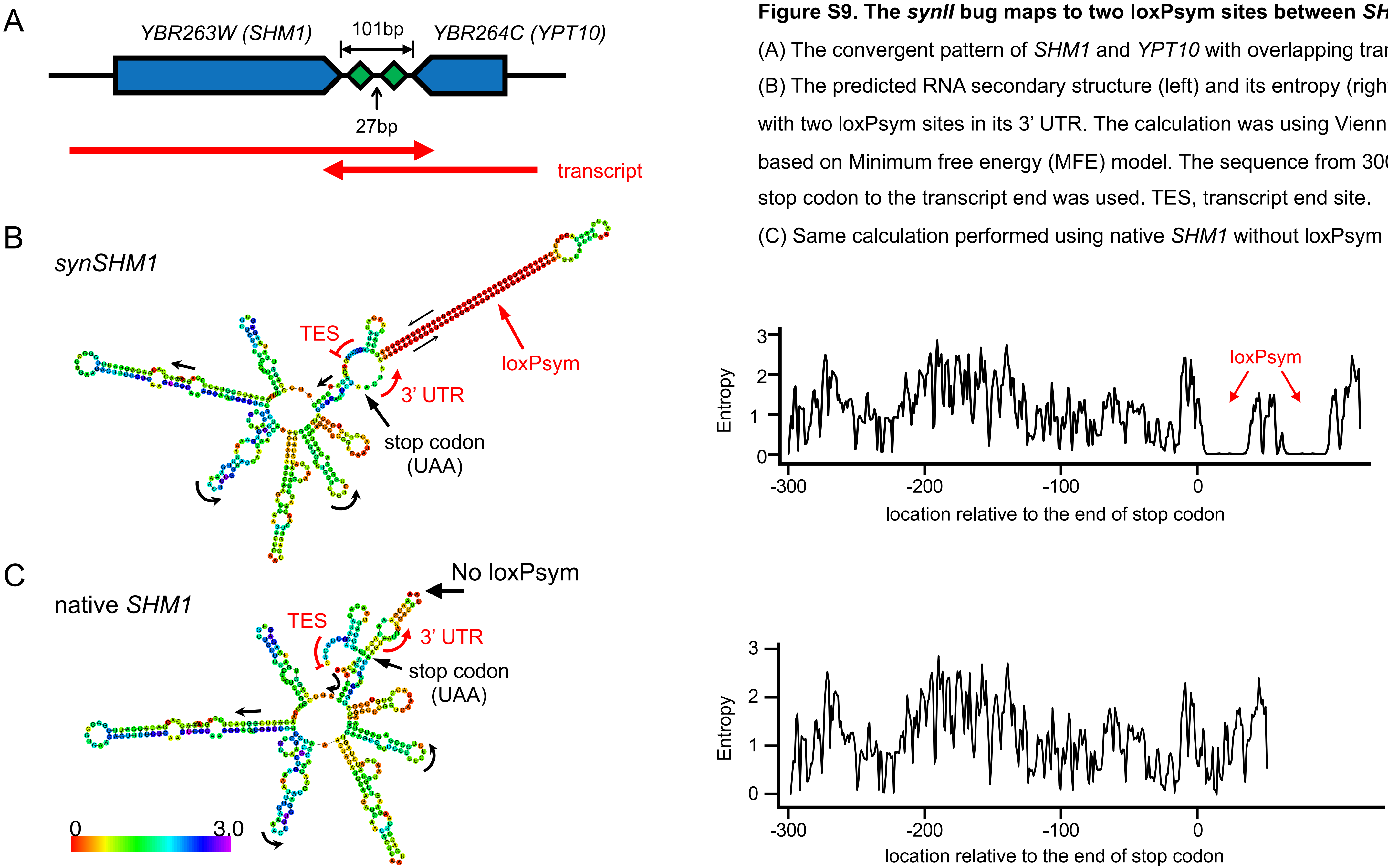


**Figure S9. The *synII* bug maps to two loxPsym sites between *SHM1* and *YPT10*.**

(A) The convergent pattern of *SHM1* and *YPT10* with overlapping transcript.

(B) The predicted RNA secondary structure (left) and its entropy (right) for *synSHM1* with two loxPsym sites in its 3' UTR. The calculation was using ViennaRNA Package based on Minimum free energy (MFE) model. The sequence from 300 nt upstream of stop codon to the transcript end was used. TES, transcript end site.

(C) Same calculation performed using native *SHM1* without loxPsym sites.

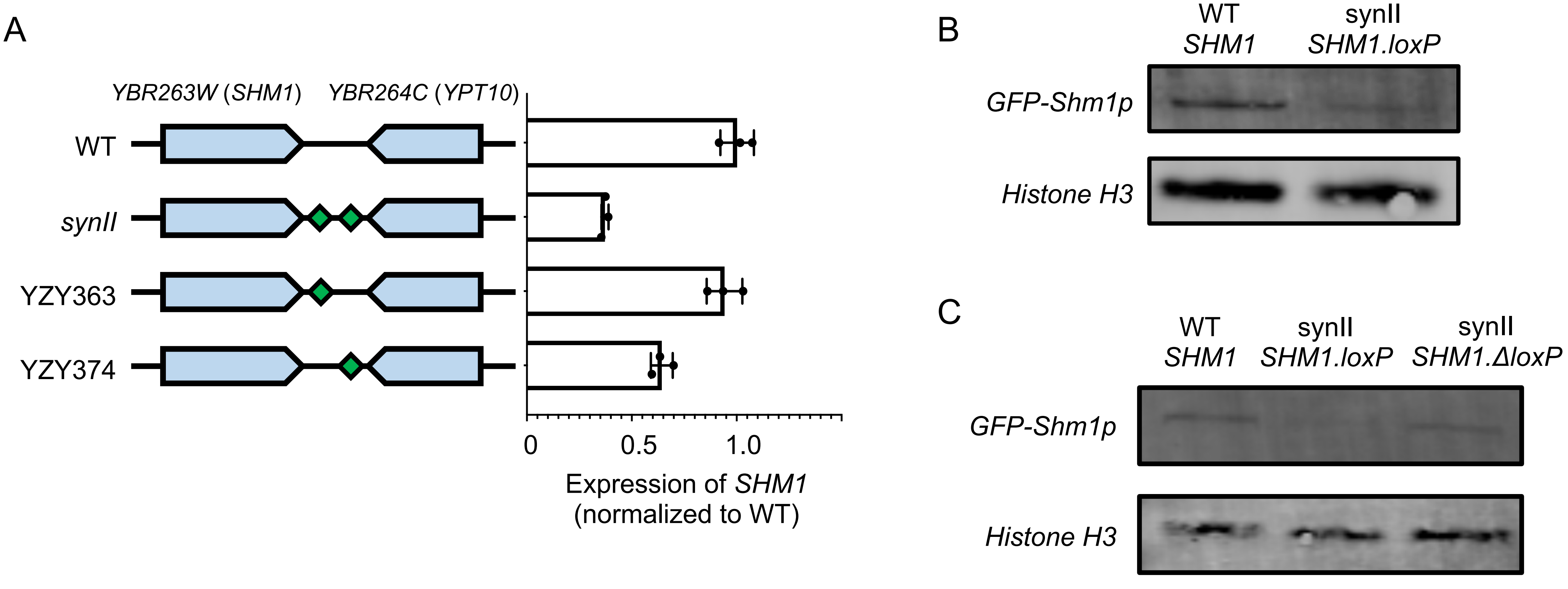


**Figure S10. The protein and transcript level of *SHM1* with or without the two loxPsym sites.**

(A) Real-time PCR to check the transcript level. YZY363, *synII* strain with only left side loxPsym site. YZY374, *synII* strain with only right side loxPsym site.

(B) Western blot to check level of N-terminally GFP-tagged Shm1p. Histone H3 was used as the internal control.

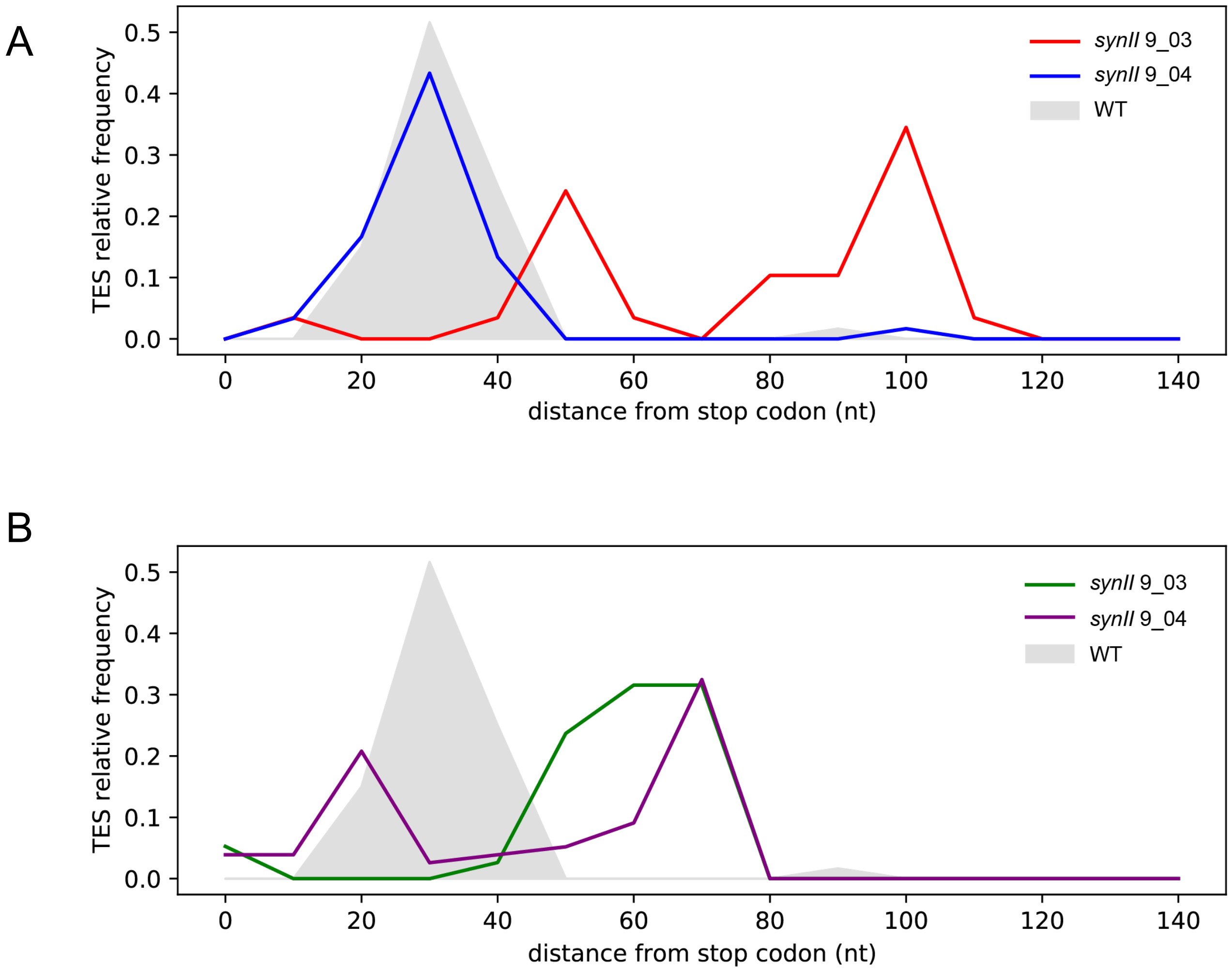


**Figure S11. Transcript end site distributions for *YPT10* (*YBR264C*).**

(A) The TES distributions of the *YPT10* transcript, in original *synII* with two loxPsym sites (9_03) and updated version with both loxPsym sites deleted (9_04).

(B) The same measurements for the *YPT10* transcript with either of loxPsym sites, in the *synII* of YZY364 and YZY374, also shown in Figure 3D.

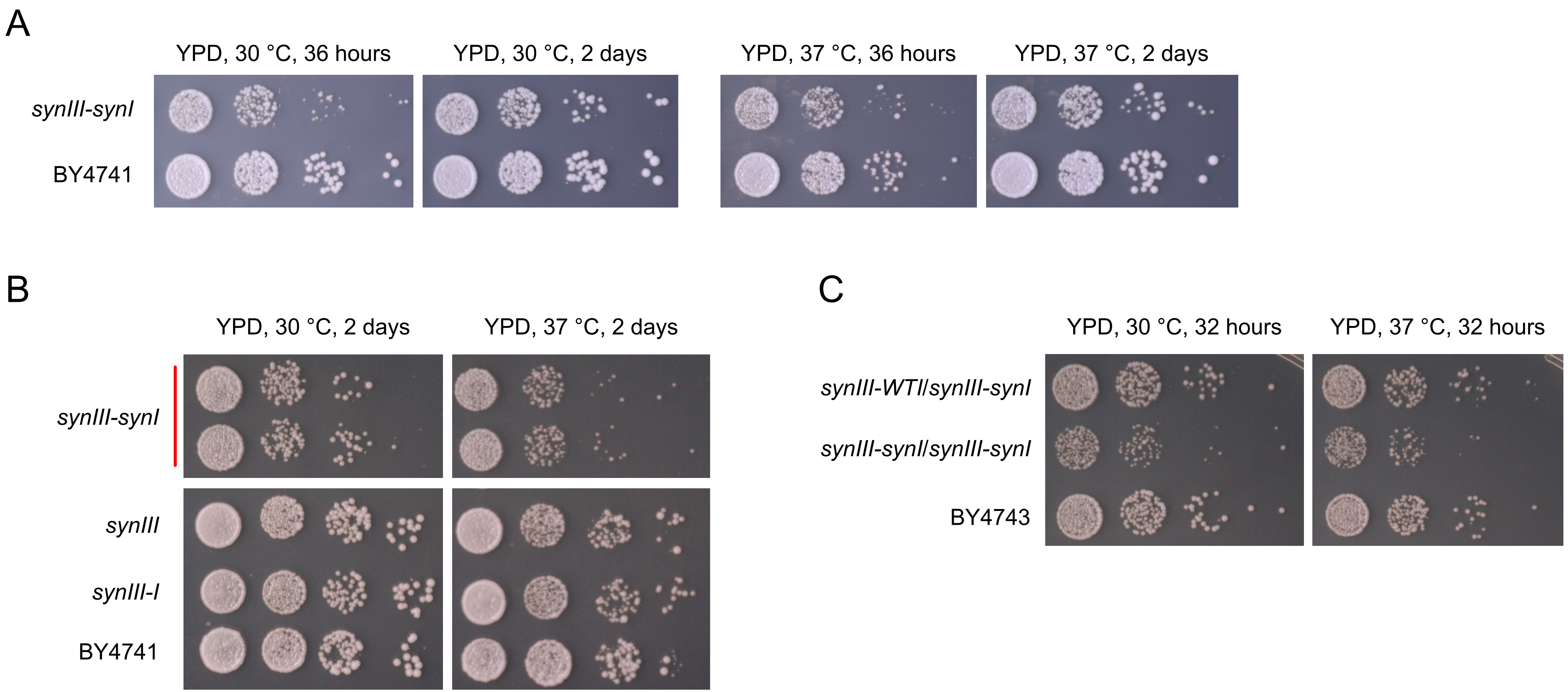


**Figure S12. Fitness assay of strains containing *synI* fused with *synIII*.**

(A) The fitness assay showing growth of serially diluted draft *synIII-synI* strain yJL671 on YPD at 30°C and 37°C.

(B) The fitness assay to check the effect of *synIII* and chromosome fusion. *synIII-synI*, the same strain as in (A). *synIII*, the strain containing separate *synIII* and wild-type chr *I*. *synIII-I*, the strain containing *synIII* fused with wild-type chr *I*.

(C) The fitness assay performed on diploids to show that the defect is recessive.

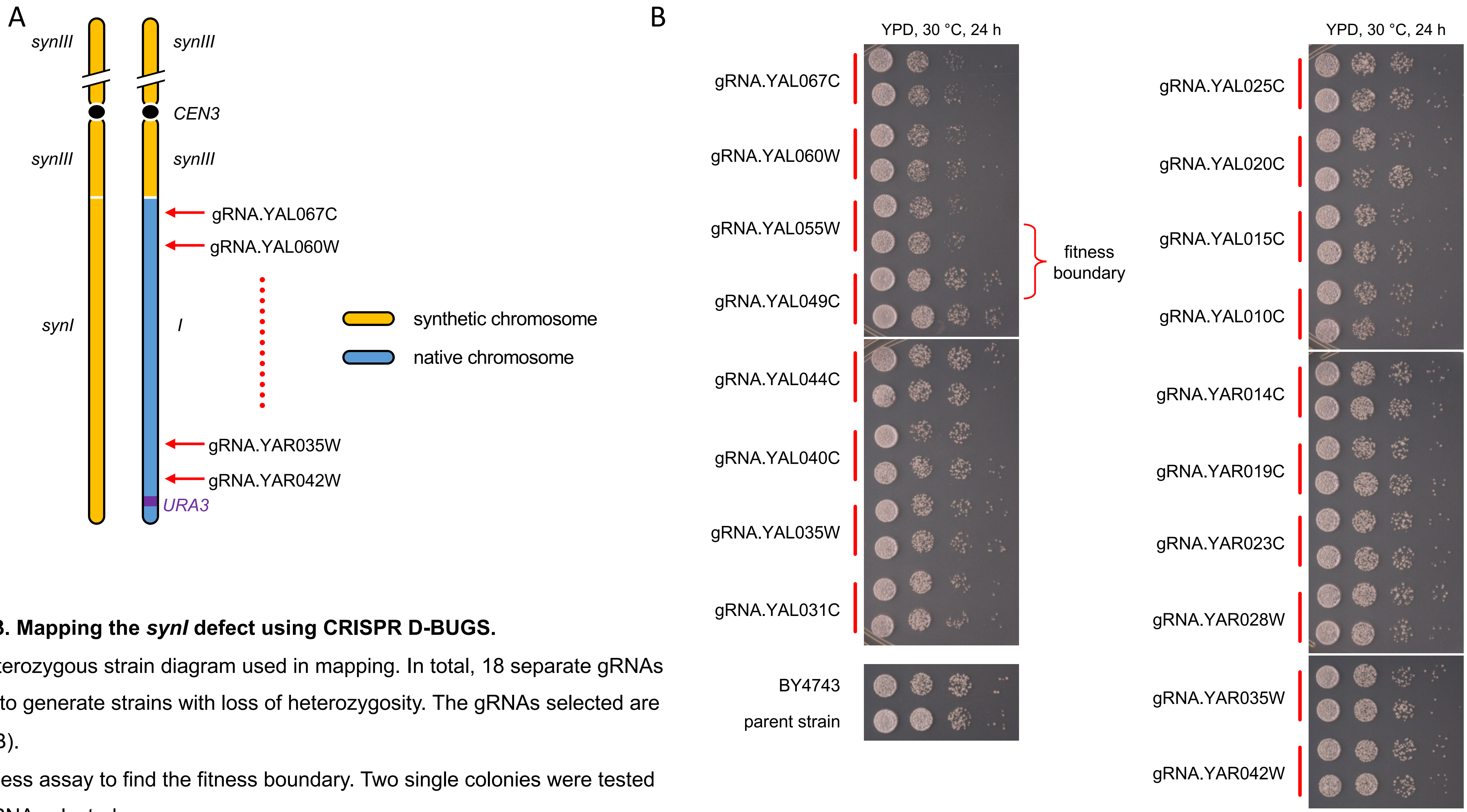


**Figure S13. Mapping the *synI* defect using CRISPR D-BUGS.**

(A) The heterozygous strain diagram used in mapping. In total, 18 separate gRNAs were used to generate strains with loss of heterozygosity. The gRNAs selected are shown in (B).

(B) The fitness assay to find the fitness boundary. Two single colonies were tested for each gRNA selected.

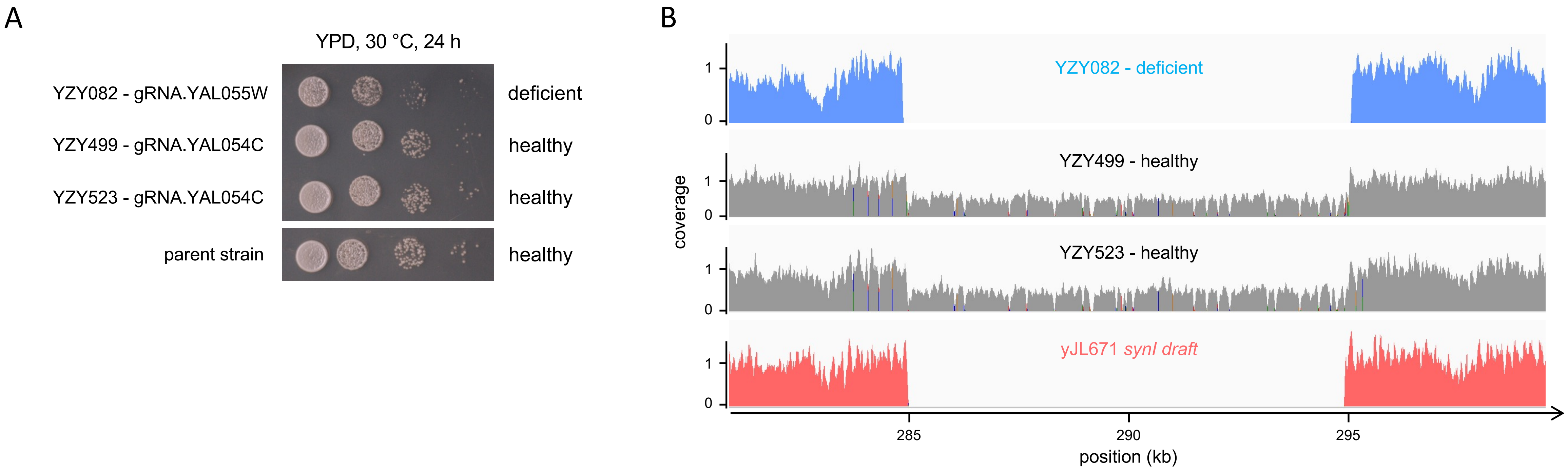


**Figure S14. The *synI* defect was mapped to a regional ~10kb deletion.**

(A) The spot assay to check the fitness of strains generating using gRNA.YAL055W (YZY082) and gRNA.YAL054C (YZY499 and YZY523). Parent strain, the diploid strain with heterozygous chr *I*.

(B) Coverage of *synI* reference genome from whole genome sequencing of strains from (A). YZY082 has no coverage while YZY499 and YZY523 only have half coverage and misaligned genome feature, indicating one allele is wild-type. Y-axis, the coverage normalized to the average genomic coverage level. X-axis, the position starting from the left end of *synIII-synI* fusion chromosome.

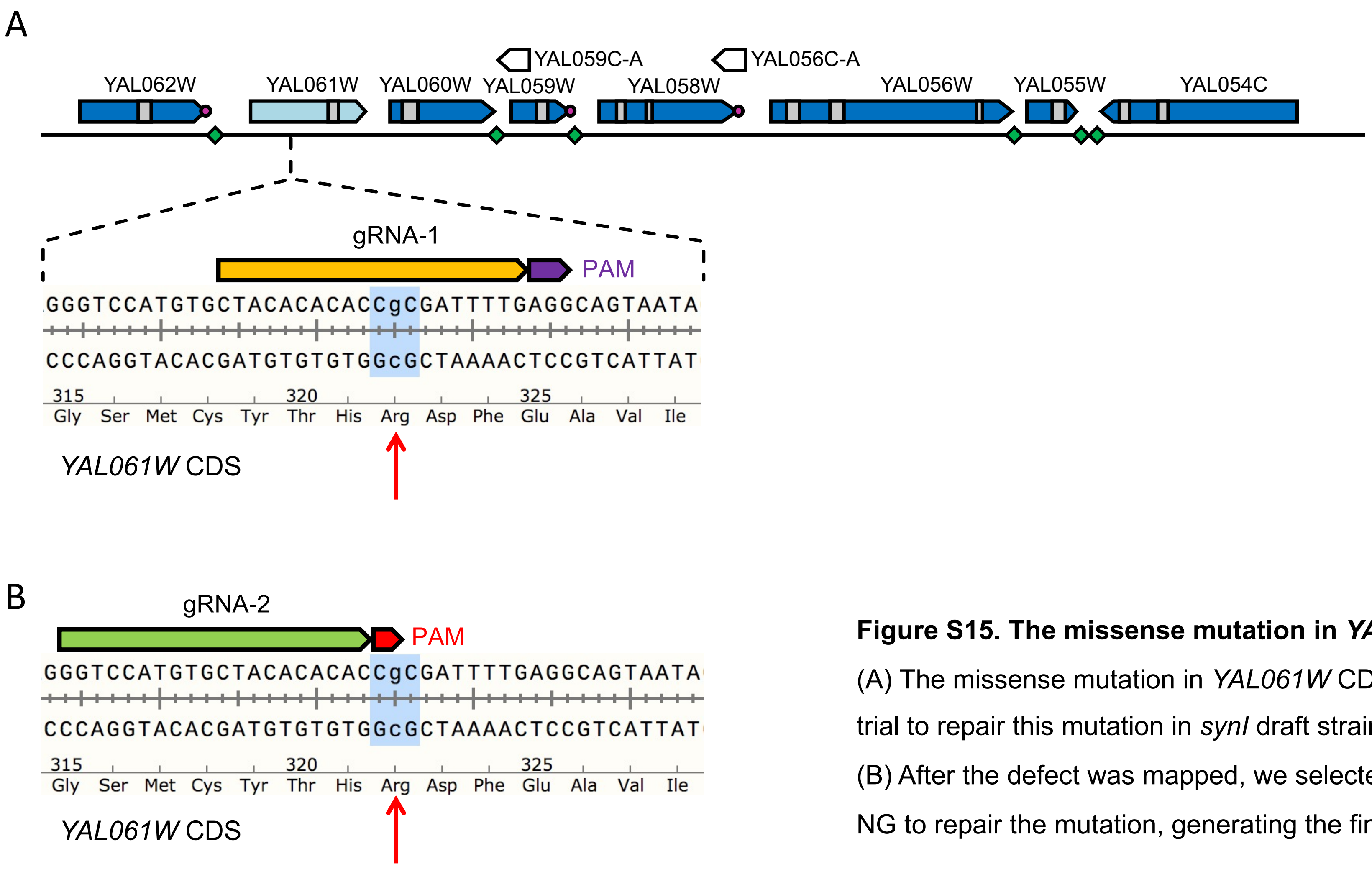


**Figure S15. The missense mutation in *YAL061W* and its repair.**

(A) The missense mutation in *YAL061W* CDS (red arrow) and the gRNA-1 used in the first trial to repair this mutation in *synI* draft strain.

(B) After the defect was mapped, we selected a new gRNA-2, compatible with SpCas9-NG to repair the mutation, generating the final and healthy *synI* strain yCTC002.

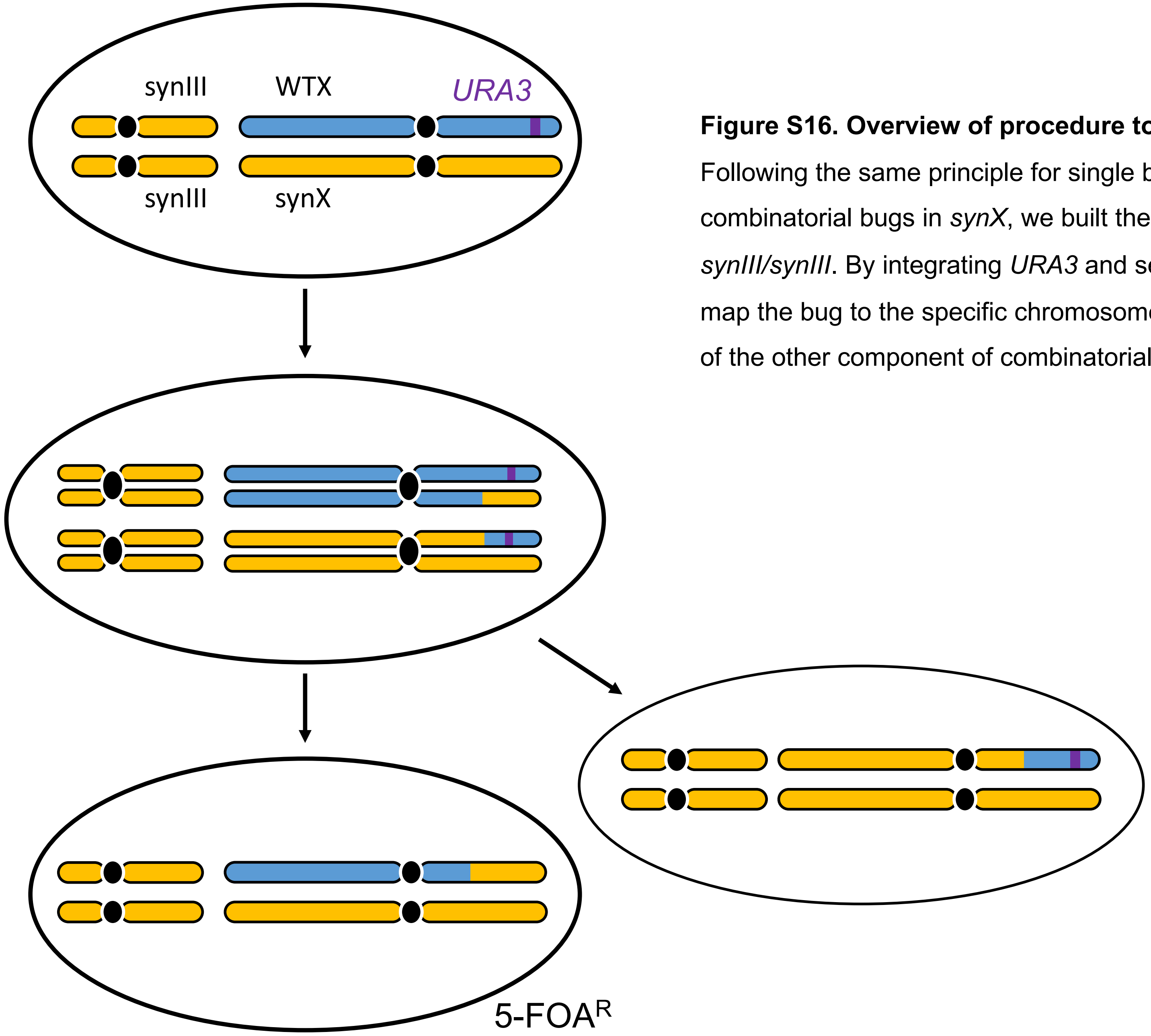


**Figure S16. Overview of procedure to map combinatorial defect using CRISPR D-BUGS.** Following the same principle for single bugs (Figure 2), if we want to map the component of combinatorial bugs in *synX*, we built the strain of heterozygous *synX/chrX*$^{+}$ but homozygous *synIII/synIII*. By integrating *URA3* and selecting the gRNA targeting at left or right arm, we can first map the bug to the specific chromosome arm at first. The same principle applies to the mapping of the other component of combinatorial bugs in *synIII*.

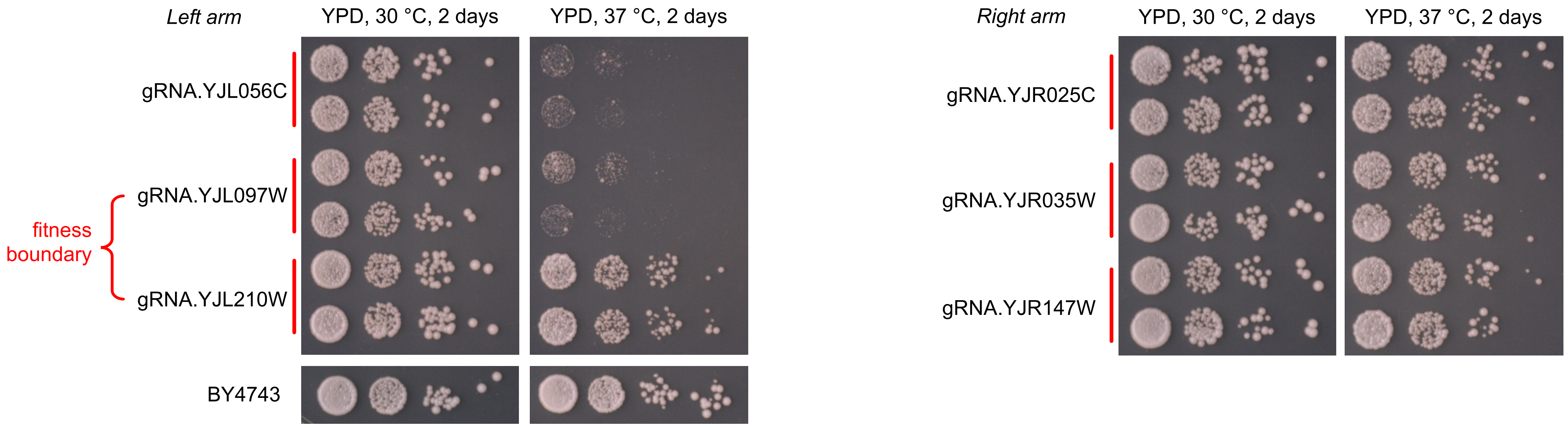


**Figure S17. First round of bug mapping for *synX*.**

We first selected 6 spatial distributed gRNAs targeting the left arm (left) and right arm (right) for CRISPR D-BUGS. The fitness assay shows that the bug is located in the left arm and we found the fitness boundary between *YJL097W* and *YJL210W*.

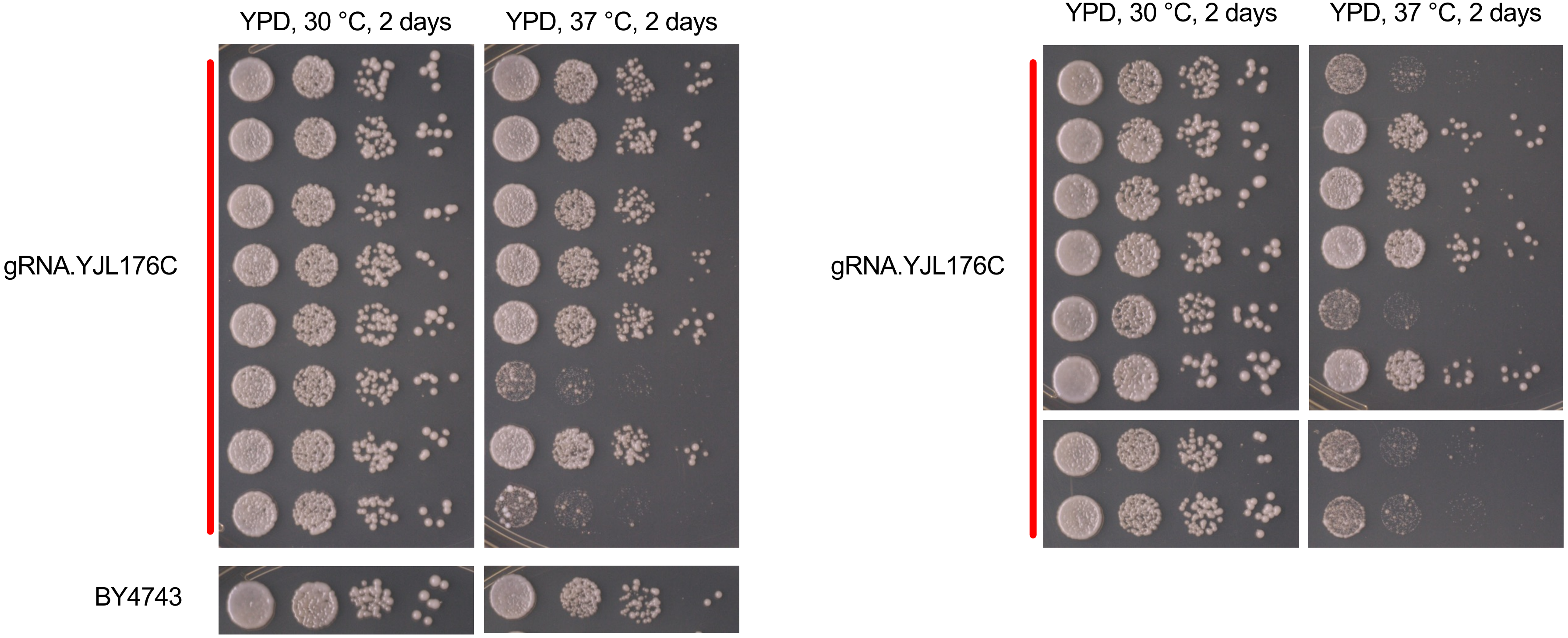


**Figure S18. Fine mapping using one single gRNA targeting at wild-type *YJL176C*.**

Individual colonies were generated using the same gRNA.YJL176C but showed a diverse level of fitness. These colonies were whole-genome sequenced and aligned to *synX* as the reference to map their recombination sites as in Figure 4B.

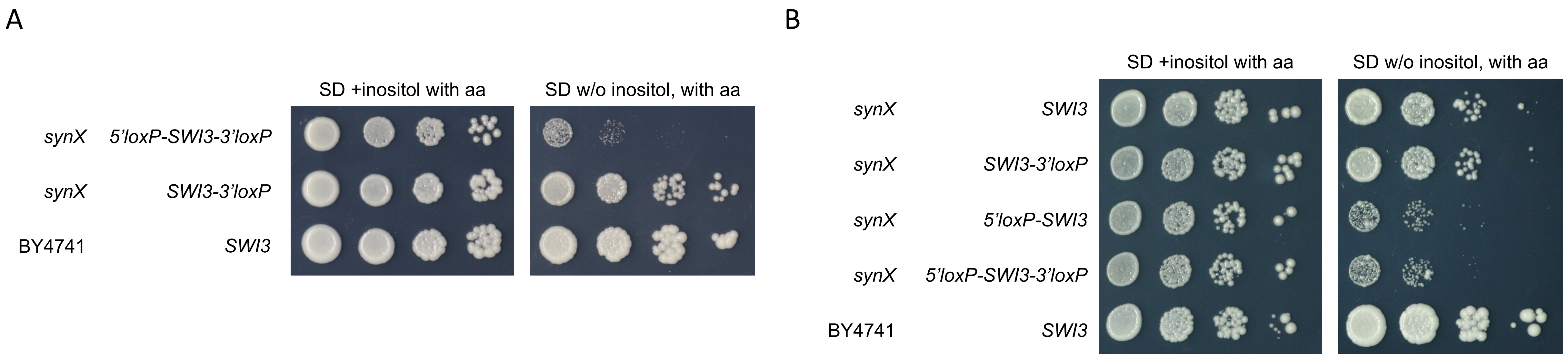


**Figure S19. Spot assay to check the inositol auxotrophy.**

(A) Growth of the original *synX* strain (chr10_9_01) with *5'loxP-SWI3.3'loxP* compared to the *SWI3.3'loxP* strain in which the loxPsym site in 5' UTR of *SWI3* was deleted. The BY4741 with wild-type *SWI3* was used as a control. The plates were incubated for 3 days at 30 °C.

(B) Yeast strains with original *synSWI3* in synX (5'loxP-SWI3-3'loxP), compared to the strain with 3'loxP deleted (*5'loxP-SWI3*), 5'loxP deleted (*SWI3-3'loxP*) or both deleted (*SWI3*).

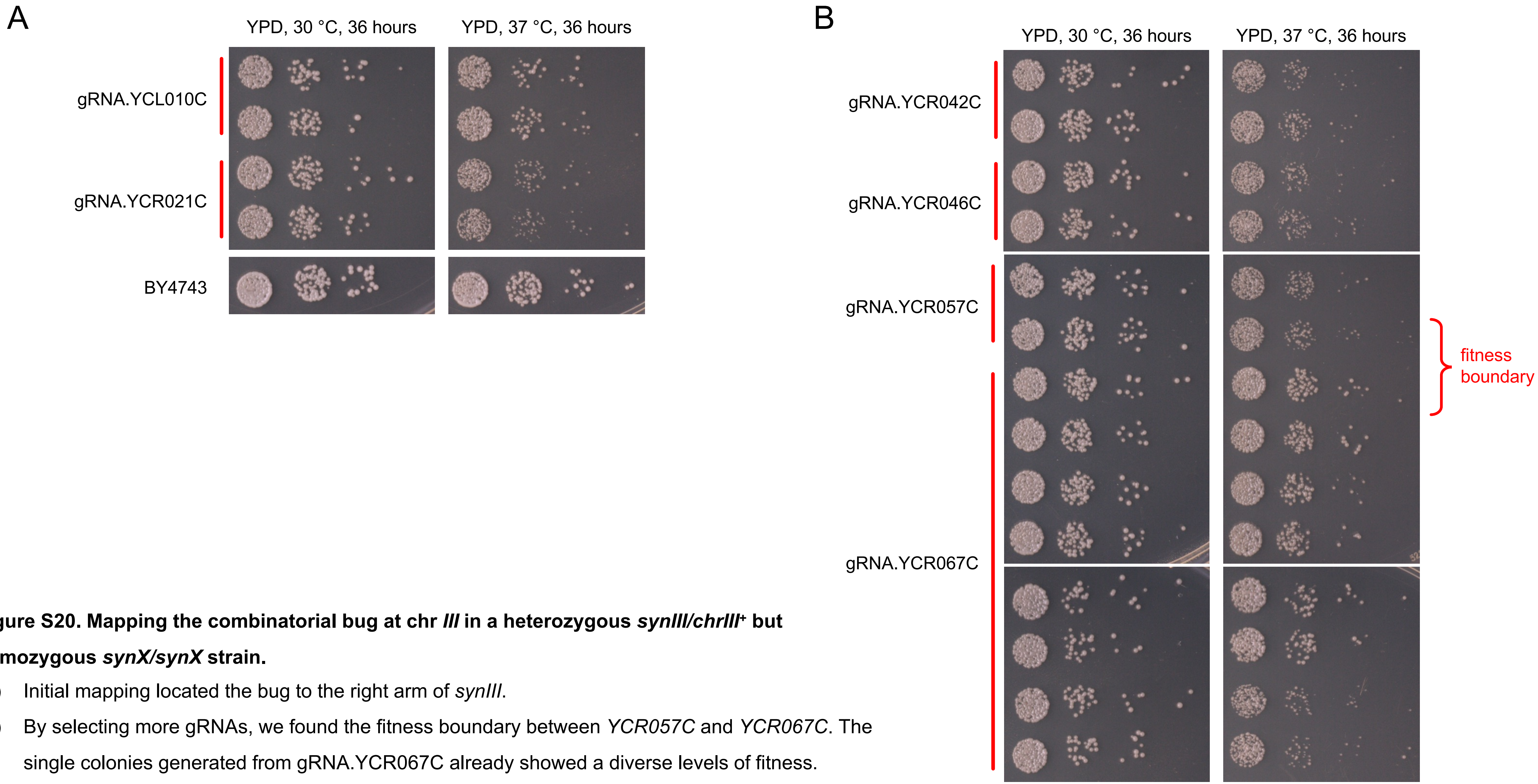


**Figure S20. Mapping the combinatorial bug at chr *III* in a heterozygous *synIII/chrIII*⁺ but homozygous *synX/synX* strain.**

(A) Initial mapping located the bug to the right arm of *synIII*.

(B) By selecting more gRNAs, we found the fitness boundary between *YCR057C* and *YCR067C*. The single colonies generated from gRNA.YCR067C already showed a diverse levels of fitness.

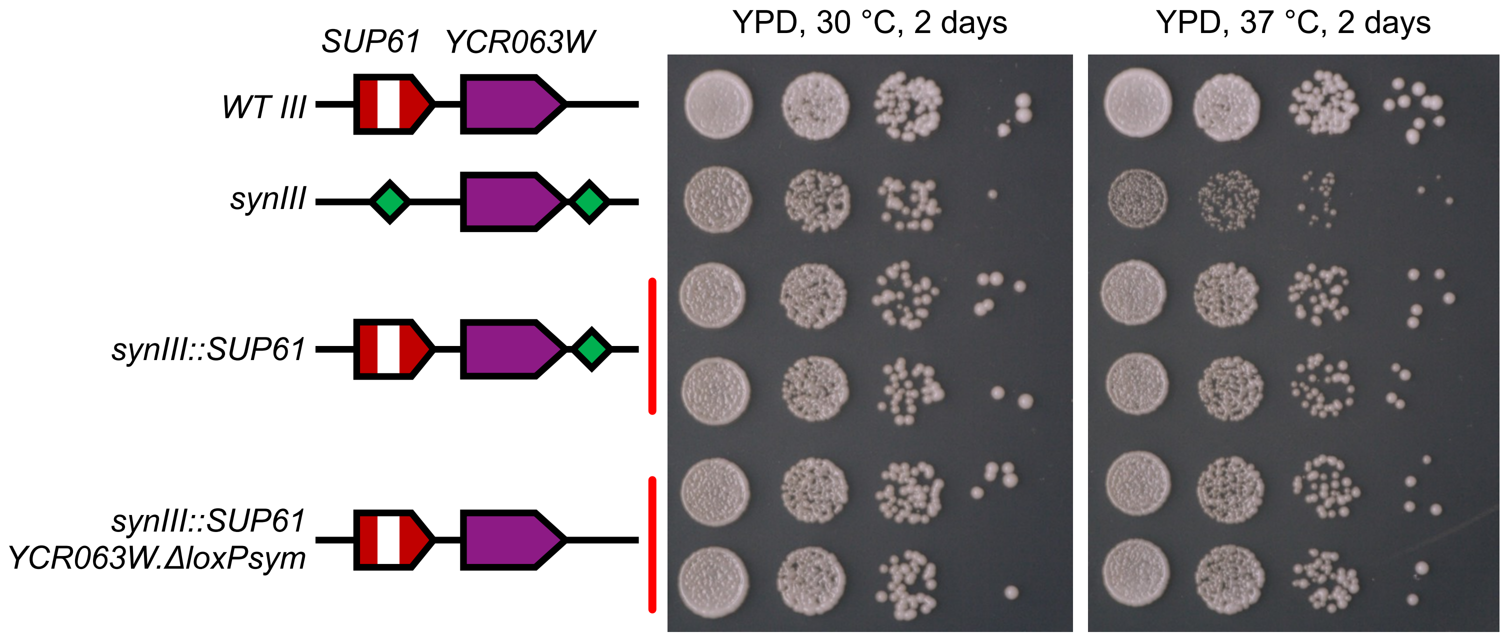


**Figure S21. Rescuing the growth defect in *synIII* and *synX* strain by integrating *SUP61* and/or deleting the right side loxPsym site.** CRISPR D-BUGS already mapped the bug to two loxPsym site (green diamond) in *synIII*. The left one was the landmark for *SUP61* deletion and the right one was integrated downstream of *YCR063W*. The integration of *SUP61* (*synIII::SUP61*) was sufficient to rescue the defect.

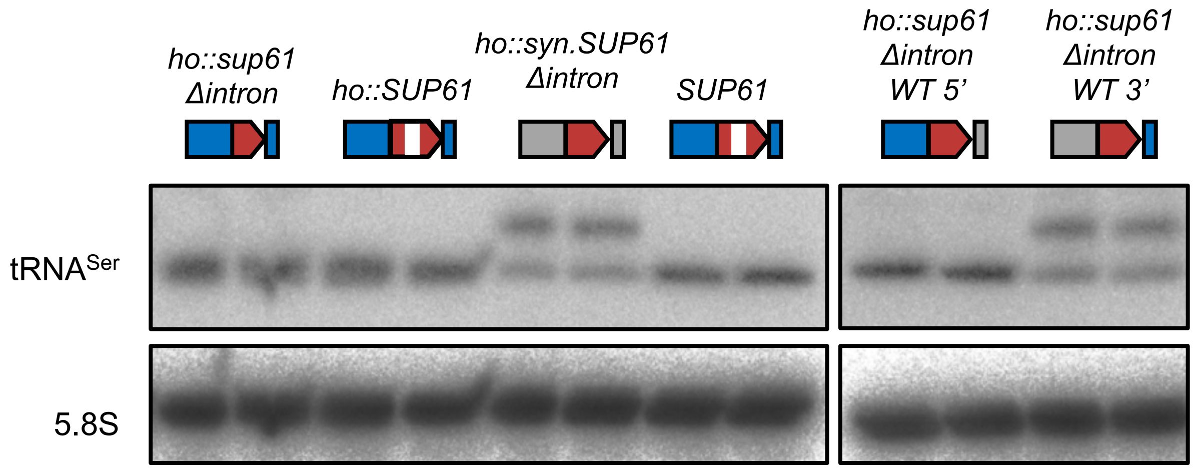


**Figure S22. Northern blot to check the quality and level of tRNA$_{Ser}$$^{CGA}$.**

The tRNA$_{Ser}$$^{CGA}$ species was expressed from different version of *syn.SUP61*. Two single colonies were tested in each group as replicates. Red, *SUP61* coding sequence. Gray, flanking sequences from *Eremothecium (Ashbya) gossypii*. Blue, the native sequences from wild-type *S. cerevisiae*. White, intron in *SUP61*.

A

| | codon | fraction(%) | tRNA gene |
|---|---|---|---|
| Ser | AGU | 0.16 | 0 |
| Ser | AGC | 0.11 | 2 |
| Ser | UCU | 0.26 | 11 |
| Ser | UCA | 0.21 | 3 |
| Ser | UCC | 0.16 | 0 |
| Ser | UCG | 0.10 | 1* |

**S288C genome**

B

| | codon | number | fraction(%) |
|---|---|---|---|
| Ser | AGU | 24 | 0.30 |
| Ser | AGC | 12 | 0.15 |
| Ser | UCU | 12 | 0.15 |
| Ser | UCA | 11 | 0.14 |
| Ser | UCC | 9 | 0.11 |
| Ser | UCG | 12 | 0.15 |

***synSWI3***

C

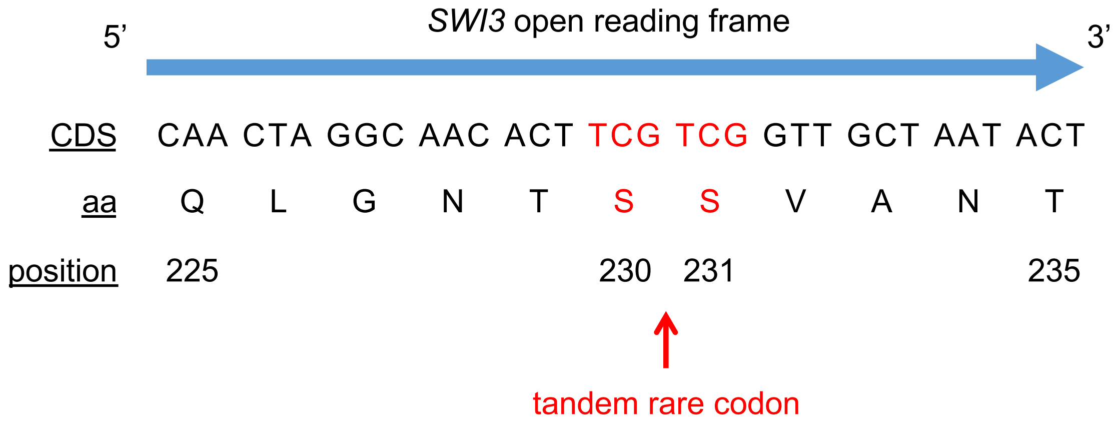


**Figure S23. Codon usage table for serine in *S. cerevisiae* genome and *SWI3*.**

(A) The codon usage for serine in wild-type yeast genome. * UCG is a rare codon that is decoded by the tRNA species encoded by *SUP61*, which is essential.

(B) The codon usage for serine in *SWI3* at synX, with tandem rare codons shown in (C). Position is the amino acid counted from the start codon.

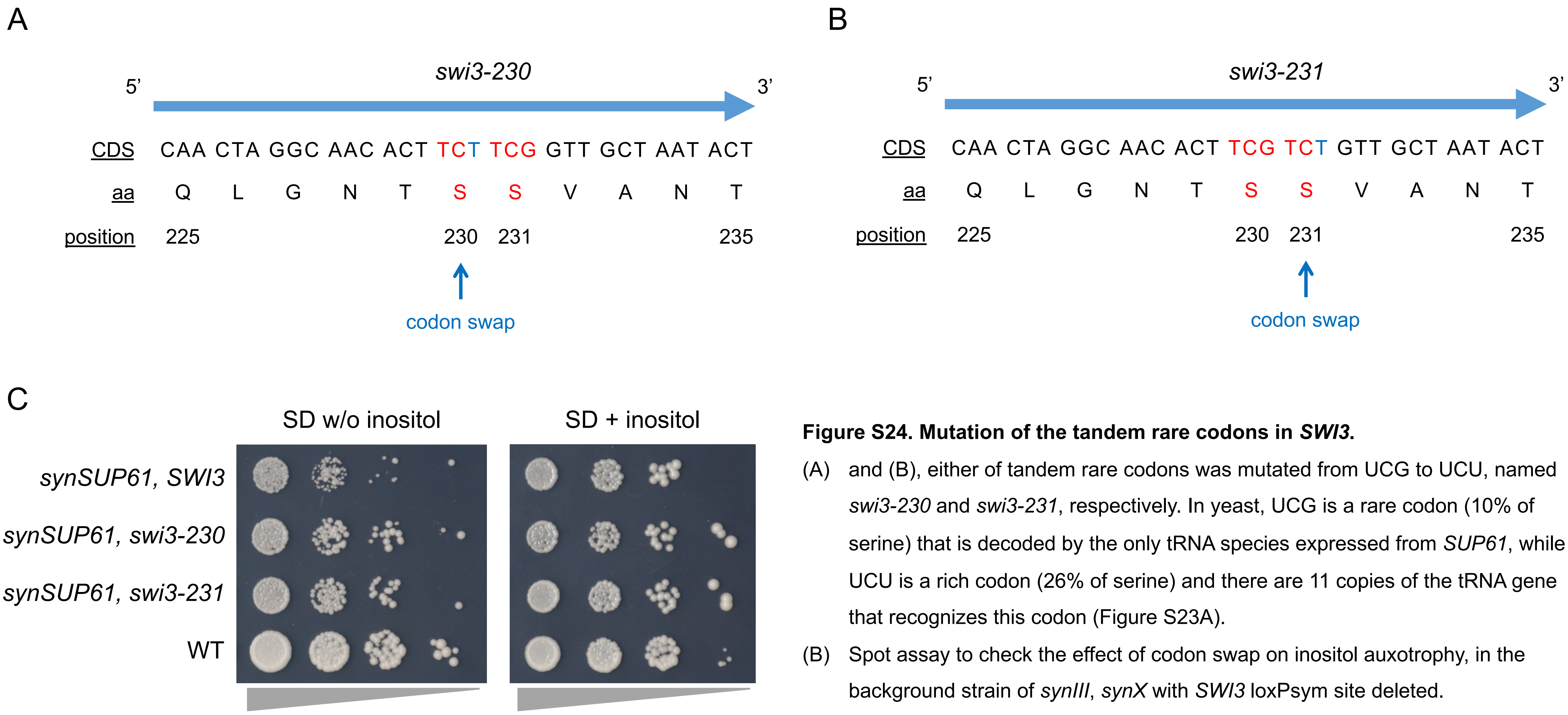


**Figure S24. Mutation of the tandem rare codons in *SWI3*.**

(A) and (B), either of tandem rare codons was mutated from UCG to UCU, named *swi3-230* and *swi3-231*, respectively. In yeast, UCG is a rare codon (10% of serine) that is decoded by the only tRNA species expressed from *SUP61*, while UCU is a rich codon (26% of serine) and there are 11 copies of the tRNA gene that recognizes this codon (Figure S23A).

(B) Spot assay to check the effect of codon swap on inositol auxotrophy, in the background strain of *synIII*, *synX* with *SWI3* loxPsym site deleted.

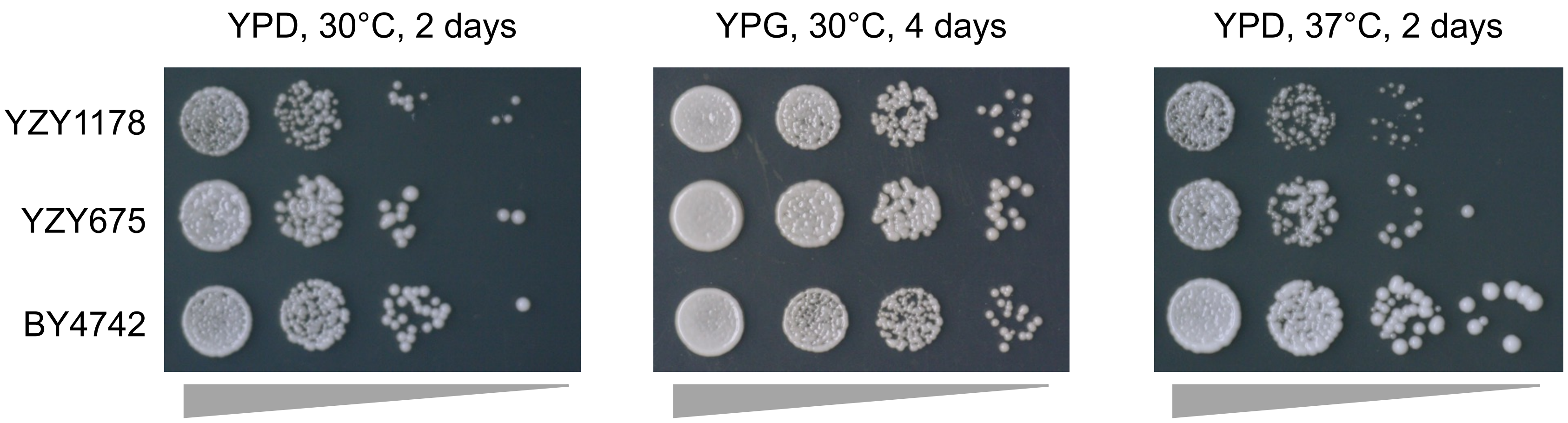


**Figure S25. Fitness assay for final strains with multiple synthetic chromosomes.**

The draft strain containing *synII*, *synIII*, *synV*, *synVI*, *synIXR*, *synX* and *synXII* is YZY1178, which is also shown in Figure 1D. YZY675 was generated by repairing all know bugs, including the *SHM1* bug in *synII* and the combinatorial bug between *SWI3* in synX and *synSUP61* in *synIII*, generating YZY675.

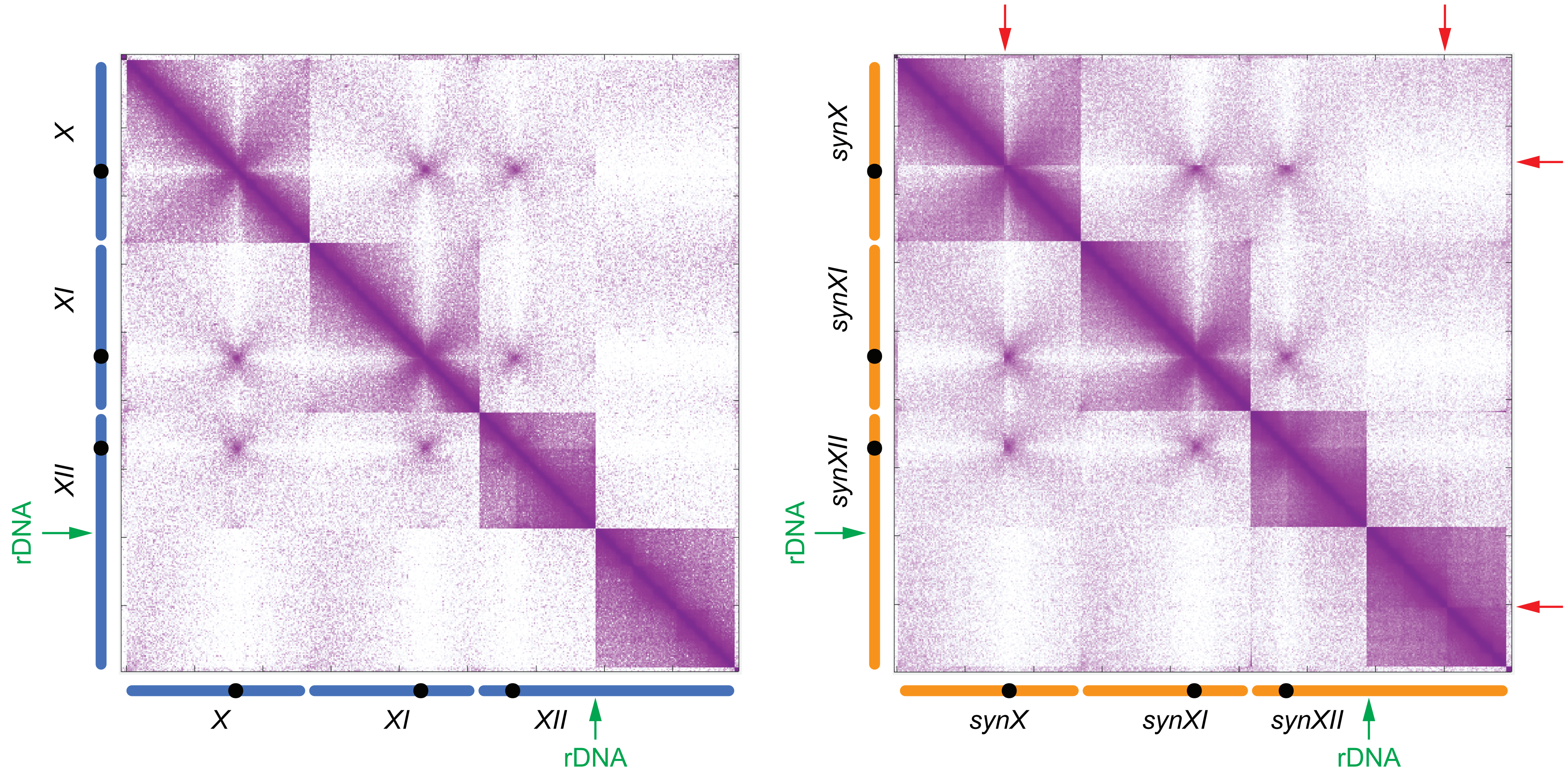


**Figure S26. Contact frequency heat map showing sharp boundaries for *synX*, *synXII*, compared with native chromosome *XI*.** The heat map shows contact frequencies from Hi-C data for native (left) and synthetic (right) chromosomes *X*, *XI* and *XII*. The boundaries at the tRNA array integration loci are highlighted with red arrows.

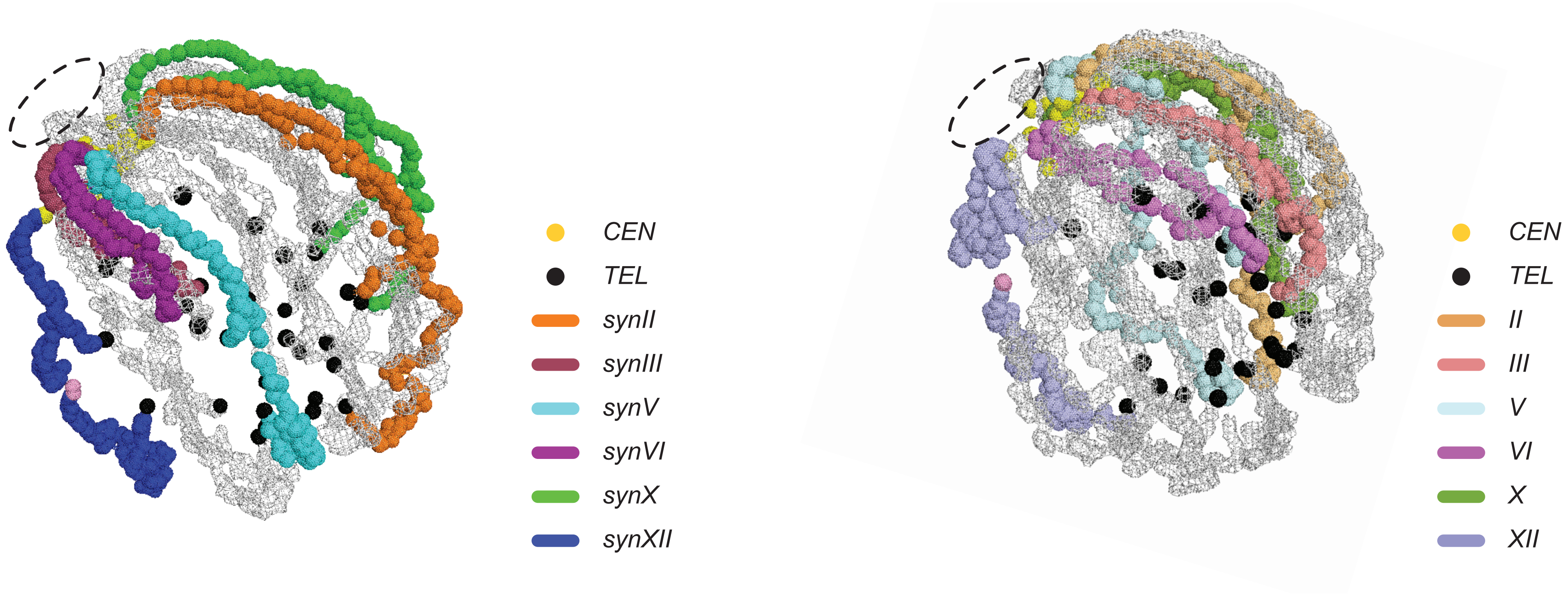


**Figure S27. The 3D chromosome trajectories of multiple synthetic chromosomes (left), compared to wild type chromosomes (right).** Gray, all other native chromosomes. The original PyMOL file for 3D chromosome organization is in Supplementary Data-1. A movie was also generated using the same labels in Supplementary Data-2.

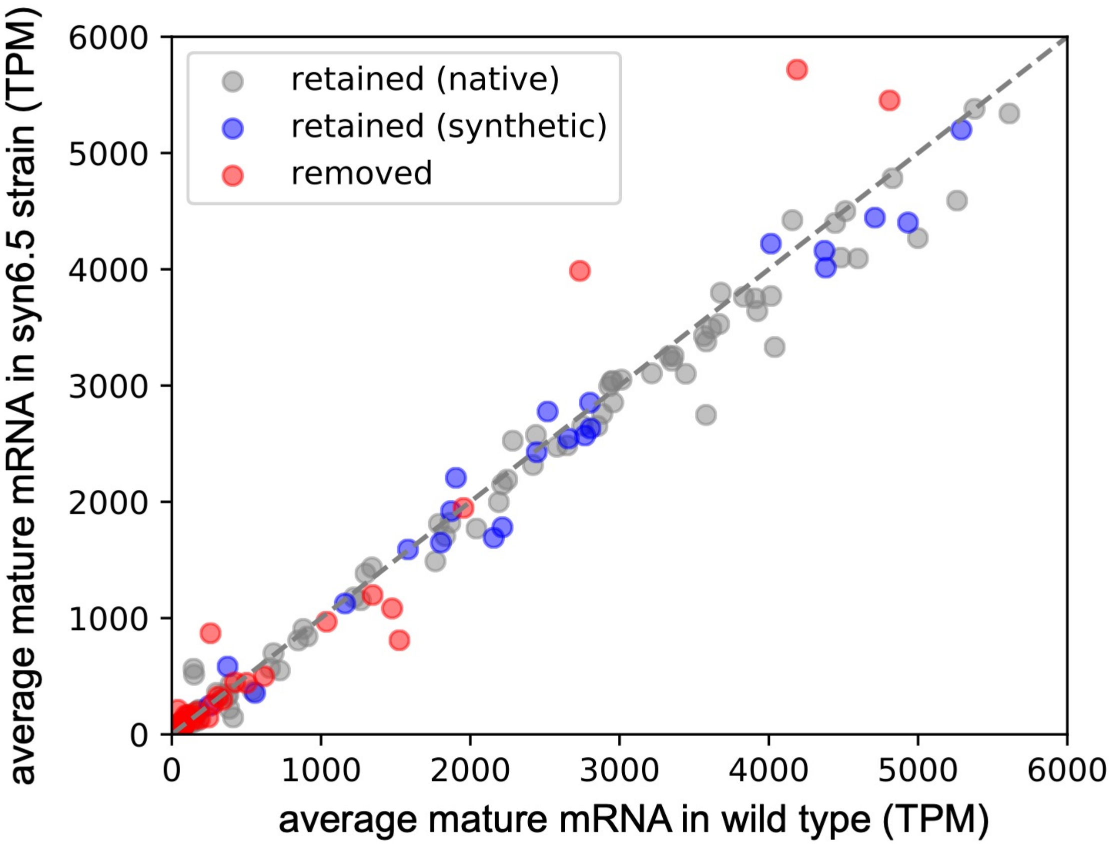


**Figure S28. Expression levels of genes in syn6.5 strain vs wild type control.** Mean salmon quantification from three replicates of Illumina stranded mRNA sequencing of the syn6.5 and wild type strains are compared for intron-containing genes. Genes that retained their introns in the syn6.5 strain on the native or synthetic chromosomes are colored in gray and blue, respectively; while genes with intron deletion are shown in red. TPM: transcripts per million.

A

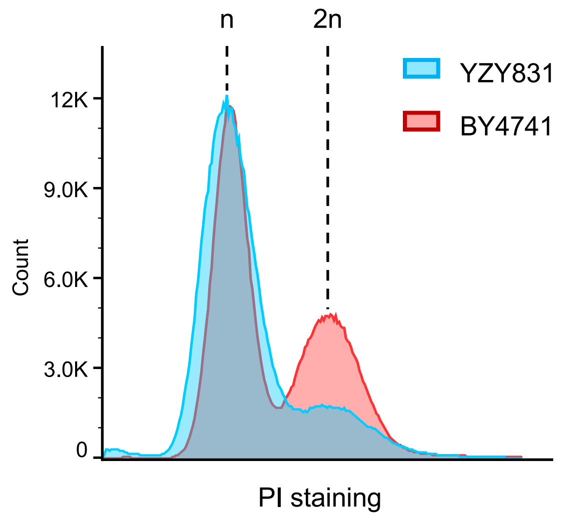


**Figure S29. Check the DNA content and sequence of the strain containing *synII*, *synIII*, *synIV*, *synV*, *synVI*, *synIXR*, *synX*, *synXII*.**

(A) Flow cytometry data confirmed the DNA content of multiple synthetic strain YZY831, with BY4741 as the wild-type haploid control.

(B) Coverage from the whole genome sequencing of YZY831, aligned to the reference sequence of multiple synthetic chromosomes.

B

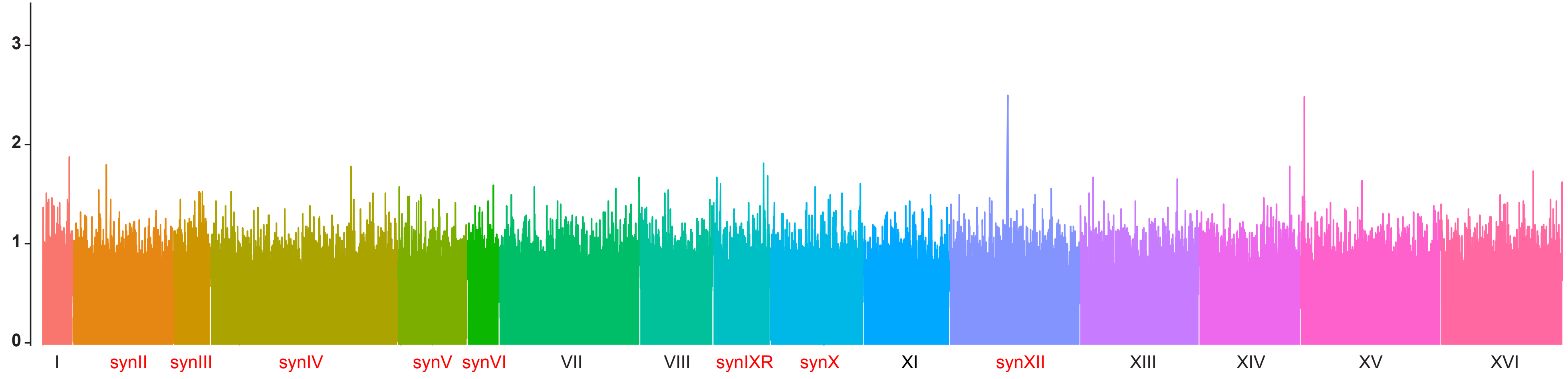

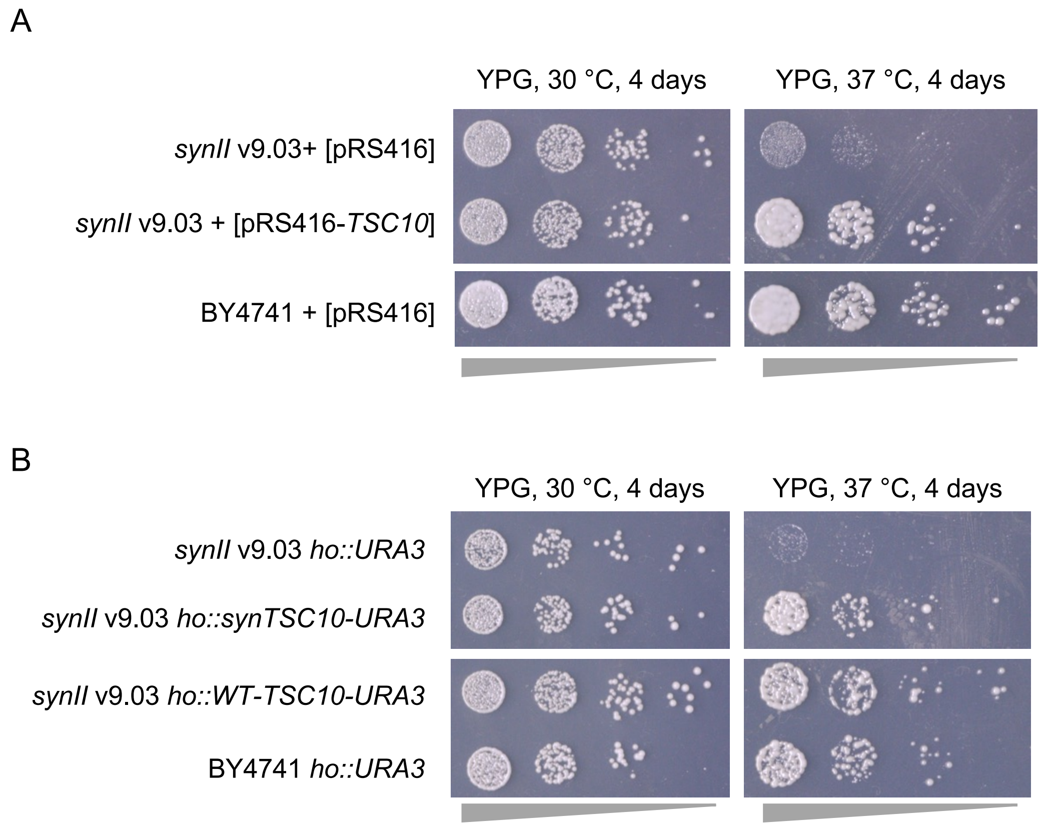


**Figure S30. Alternative method to rescue the *synII* growth defect.**

(A) Spot assay for the original *synII* strain (v9.03) transformed with empty vector (pRS416) or second copy of wild-type *TSC10* under the control of its native promoter (pRS416-TSC10). BY4741 was transformed with the empty vector as the control.

(B) Spot assay for the *synII* v9.03 with either only *URA3*, a second copy of *TSC10* with synthetic PCRtags, or a copy of *TSC10* with wild-type PCRtags integrated at the *HO* locus. BY4741 was integrated with one copy of *URA3* at the *HO* locus as a control.

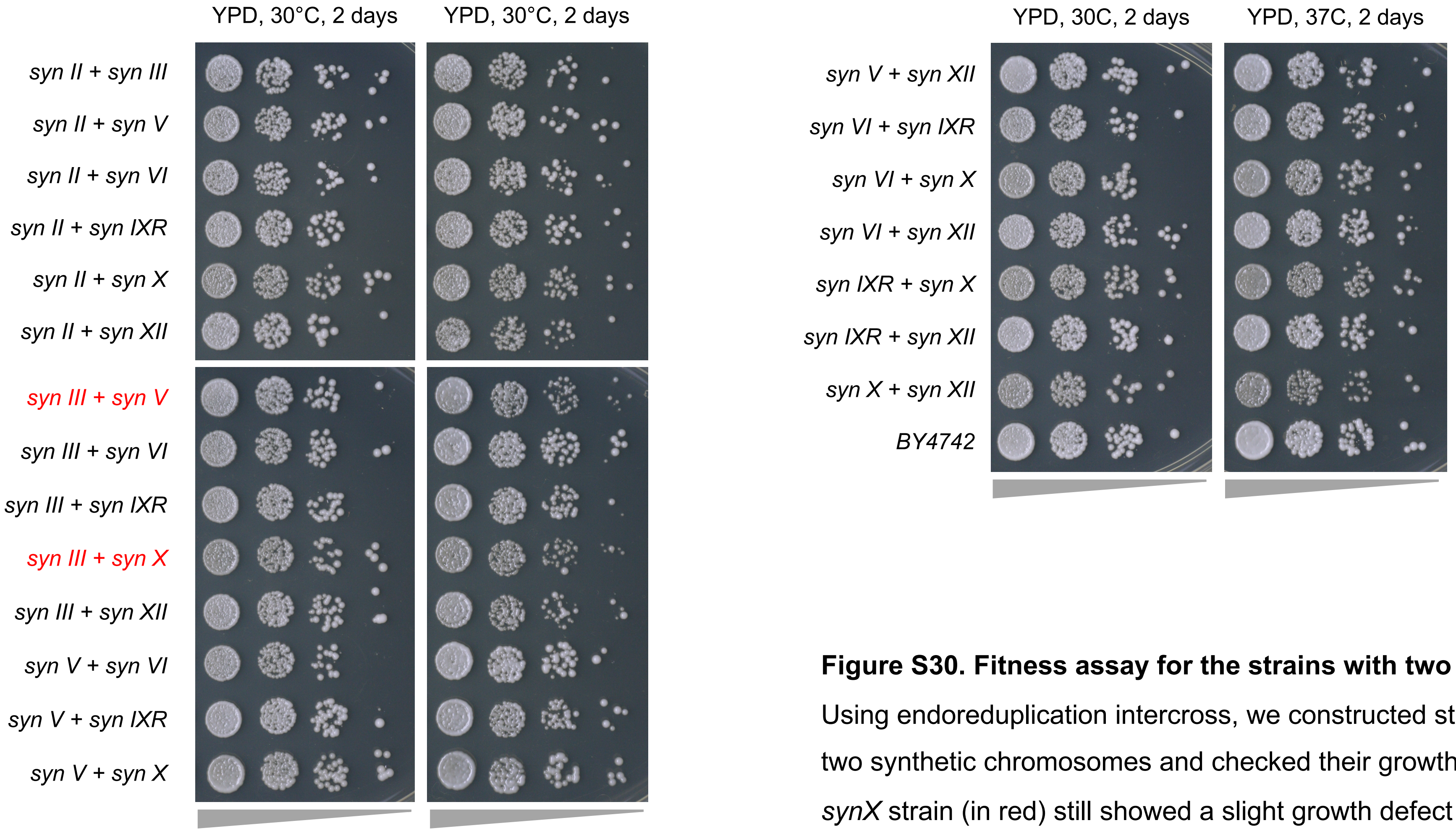


**Figure S30. Fitness assay for the strains with two synthetic chromosomes.**

Using endoreduplication intercross, we constructed strains with all possible combinations of two synthetic chromosomes and checked their growth on YPD. The *synIII, synV* and *synIII, synX* strain (in red) still showed a slight growth defect at high temperature.
